# Biofabrication of an *in situ* hypoxia-delivery scaffold for cartilage regeneration

**DOI:** 10.1101/2025.03.04.641405

**Authors:** R. Di Gesù, A. Palumbo Piccionello, G. Vitale, S. Buscemi, S. Panzavolta, M.F. Di Filippo, A. Leonarda, M. Cuccia, A. Di Prima, R. Gottardi

## Abstract

Osteoarthritis (OA) is a debilitating joint condition affecting millions of people worldwide, triggering painful chondral defects (CDs) that ultimately compromise the overarching patients’ quality of life. Currently, several reconstructive cartilage techniques (RCTs) (i.e.: Matrix-assisted Autologous Chondrocytes Implantation - MACI) has been developed to overcome the total joint replacement (TJR) limitations in the treatment of CDs. However, there is no consensus on the effectiveness of RCTs in the long term, as they do not provide adequate pro-regenerative stimuli to ensure complete CDs healing. In this study, we describe the biofabrication of an innovative scaffold capable to promote the CDs healing by delivering pro-regenerative hypoxic cues at the cellular/tissue level, to be used during RCTs. The scaffold is composed of a gelatin methacrylate (GelMA) matrix doped with hypoxic seeds of GelMA functionalized with a fluorinated oxadiazole (GelOXA), which ensures the delivery of hypoxic cues to human articular chondrocytes (hACs) embedded within the scaffold. We found that the GelMA/GelOXA scaffold preserved hACs viability, maintained their native phenotype, and significantly improved the production of type II collagen. Besides, we observed a reduction in type I and type X collagen, characteristic of unhealthy cartilage. These findings pave the way for the regeneration of healthy, hyaline-like cartilage, by delivering hypoxic cues even under normoxic conditions.

Furthermore, the GelMA/GelOXA scaffold’s ability to deliver healing signals directly to the injury site holds great potential for treating OA and related CDs, and has the potential to revolutionize the field of cartilage repair and regenerative medicine.

## INTRODUCTION

Osteoarthritis (OA) is a joint disorder with a heavy socioeconomic burden on patients and healthcare systems^1,2^. Over 250 million people worldwide are estimated to be affected by symptomatic OA, drastically limiting patients’ ability to perform daily activities^3^. Recent epidemiologic studies^3^ have underlined that the prevalence of OA has been increasing annually, with a high correlation with aging, and obesity. Despite extensive efforts to identify a treatment for OA, no disease-modifying pharmacological therapy has yet been developed^4^. In this complex scenario, the only resolutive treatment remains total joint replacement (TJR), which is the end-point approach most widely adopted^5–7^ based on the surgically removal of damaged tissues, and their replacement with artificial prostheses. Unfortunately, the failure rate associated with TJR remains relatively high, mainly due to postoperative complications such as periprosthetic infections^8^ or prosthesis misalignment^9,10^. Furthermore, prostheses have a limited lifetime of about 10 to 25 years, and often require revision surgery^11,12^, making TJR challenging in young patients with post-traumatic OA. This is highly relevant as up to 36% of athletes suffer from full-thickness focal chondral defects^13^, and up to 89% of NBA players and 20% of american football players have articular cartilage abnormalities^14^. Recently, different surgical alternatives to TJR have been developed, and are often adopted as the first-line approach for early-stage OA. Among these alternatives, reparative cartilage techniques (RCT), such as microfracture (MF)^15,16^, autologous chondrocyte implantation (ACI)^17,18^, and matrix-assisted autologous chondrocyte implantation (MACI)^19^, are the most representative. The main rationale for RCTs is to induce a pro-regenerative response that stimulates the formation of new functional cartilage^20^. Unfortunately, although clinical outcomes ^19,21–23^ report RCTs as promising alternatives to TJR, their success rate in the long term is relatively low, as only 57% of patients show complete recovery after treatment^19^. This is mainly due to a different biochemical composition of the new formed cartilage, which is composed of fibrocartilaginous fibers rich in collagen I^24,25^. This kind of cartilage, is less resistant to mechanical solicitations compared to the native hyaline cartilage (HC)^26^ that is mainly composed a dense extracellular matrix (ECM) rich in glycosaminoglycans (GAGs), proteoglycans, and type II collagen^27,28^. The HC composition is completed by the presence of ECM-embedded chondrocytes characterized (articular chondrocytes – ACs) by low metabolism and an overall limited healing capacity^29^. For this reason, the molecular pathways involved in chondrocyte biology are of great interest^30,31^ to shed light on the poor regenerative capacity of ACs that drastically limits the self/repair of hyaline cartilage. Several molecular mechanisms have been considered, including autophagy-dependent pathways^32^ and hypoxia-dependent^33^ pathways. Interestingly, the latter pathway has been reported to be of preponderant in regulating the phenotype and fate of ACs. It is well established that articular cartilage is constitutively hypoxic, and the metabolism of ACs has evolved to work optimally in an environment with 1–10% O_2_^34^. Such a hypoxic environment was reproduced during different studies *in vitro*^35–39^ to improve our understanding of ACs biology. Interestingly, all of these studies founded that low oxygen tension contributes to the maintenance of the native ACs phenotype, promoting the expression of pro-anabolic chondral genes such as the type II collagen, which is the hallmark of HC. Consequently, tissue engineering strategies that leverage the control of oxygen concentration to promote cartilage regeneration have been proposed^40^. Recently, this approach has been applied to treat osteochondral and chondral lesions, by fabricating scaffolds able to modulate oxygen concentration at the site of implantation. The common paradigm was the addition of oxygen retaining agents like hemoglobin^41^, myoglobin^42^, dioxides^43^, or peroxides^44^ into the scaffolds. However, despite the advanced technology of these approaches, the full extent to which controlled oxygen release may affect type II collagen production has not been fully elucidated. In addition, oxidative stress induced by dioxides, and peroxides, can drastically reduce cell viability and proliferation. Consequently, several studies have reported a different approach relying on the fabrication of 3D supports enriched with molecules such as oxygen scavengers, which can lower the oxygen concentration^45^. However, several drawbacks have been observed after the use of these classes of compounds, such as a low bioavailability, and a toxicity of their metabolites^46^. In this study, we aim to fill the gap between the deep knowledges on the pro-chondrogenic effect hypoxia-driven, widely studied and demonstrated *in vitro*, and the practical exploitation of those technologies to overcome the striking effects of chondral lesions. Our work intends to overcomes the technical restraints that still hamper the application of hypoxic cues *in situ* to treat chondral lesions, limiting the high potential of this technology. The technology we developed here leverages the pro-chondrogenic effects of hypoxia while developing a translationally feasible scaffold-based approach for restoring chondral lesions. Specifically, we describe the development of a hypoxic scaffold as an active matrix for RCTs, cellularized with human articular chondrocytes (hACs). Such a matrix is made of gelatin methacrylate (GelMA), enriched with GelMA chemically functionalized with biocompatible fluorinated oxadiazoles^47,48^, (GelMA-GelOXA), whose oxygen sequestering properties have been previously reported^49^. We found that our scaffold is capable to reduce the oxygen concentration in a controlled manner, creating an optimal hypoxic microenvironment for hACs. Our hypothesis is that such a hypoxic microenvironment better preserves the native chondrocyte phenotype, promoting the deposition of new ECM rich in type II collagen, ultimately enabling a functional cartilage regeneration. Although our results confirm our hypothesis, more advanced *in vivo* studies will be designed and conducted to make stronger our work.

## METHODS

### Materials

Solvents and reagents were purchased from Sigma-Aldrich (reagent grade) and were not further purified. Gelatin type A (from porcine skin, Sigma Aldrich, 250 Bloom) was used.

Compound **1** was prepared as previously described^50^. Compounds **2** and **3** were prepared according to a previously reported procedure (see below)^51^. The melting points were determined using a REICHART-THERMOVAR hot-stage apparatus.

### Synthesis of 3-pentadecafluoroheptyl-5-pentafluorophenyl-1,2,4-oxadiazole (3) (OXA)

Pentadecafluoroheptylamidoxime **1** (428 mg, 1 mmol) was suspended in toluene (10 mL), then pyridine (0.09 g, 1 mmol) and pentafluorobenzoyl chloride (254 mg, 1.1 mmol) were added and the mixture was stirred at room temperature overnigth. The solvent was removed by evaporation, and the solid residue was filtered through washing with water (3x20 mL), obtaining the *O*-pentafluorobenzoyl-pentadecafluoroheptylamidoxime **2** (603 mg, 97%, mp 154–6°C [lit. 155-156]^51^. The solid was melted at 160°C for 1 h and, after cooling, was treated with hexane. The filtrate was reduced *in vacuo* to obtain 3-pentadecafluoroheptyl-5-pentafluorophenyl-1,2,4-oxadiazole **3** (453 mg, 77%): mp, 36–8°C [lit. 37-38]^51^.

### Synthesis of Gelatin-Oxadiazole (GelOXA)

Gelatin (100 mg) was dissolved in DMSO (5 mL) and triethylamine (TEA) (55 µL) was added. Oxadiazole **3** (100 mg) was dissolved in DMSO (1 mL) and added to one half. After a few minutes, the yellowish gel formed was left at room temperature overnight. Ethyl acetate (40 mL) was added to the mixture, and the resultant mixture was centrifuged (10 min, 4000 rpm) to remove the solvent and unreacted materials. The treatment was repeated 2 times. The residue was lyophilized overnight, yielding 189 mg of GelOXA.

### Characterization of Gelatin-Oxadiazole (GelOXA)

#### Magic Angle Spinning solid state Nuclear Magnetic Resonance (MAS-ssNMR)

^19^F MAS NMR spectrum was obtained at room temperature using a Bruker Avance II 400 MHz (9.4 T) spectrometer operating at 376.49 MHz for the ^19^F nucleus with an MAS rate of 25 kHz, a 90° pulse on the ^19^F of 4 µs, a repetition delay of 4 s, and 32 scans. Sample (around 50 mg) was compressed in a 2.5-mm zirconia rotor with VESPEL caps.

#### Attenuated total reflectance Fourier transform infrared (ATR-FTIR)

Fourier transform infrared (FTIR) spectra were recorded using a Thermo Scientific Nicolet iS10 FTIR spectrometer equipped with an ATR sampling device, using a Germanium crystal as the internal reflection element. The infrared spectra were acquired at room temperature in transmittance mode from 4000 to 650 cm^-1^ with a resolution of 2 cm^-1^.

#### X-Ray diffraction (XRD)

XRD patterns were recorded on powdered samples using a Philips X’Celerator diffractometer equipped with a graphite monochromator in the diffracted beam. CuKα radiation (40 mA, 40 kV, 1.54 Å) was used. The 2θ range investigated was from 3° to 50° with a step size of 0.2° and a time/step of 5s.

#### Thermogravimetric analysis (TGA)

Thermogravimetric analysis (TGA) was performed using SDT Q600 (TA Instruments). Heating was performed in a platinum crucible under an air flow (100 mL/min) at a rate of 3.50 °C/min to 120.00 °C and then at 10.00 °C/min to 800.00 °C. Sample weights were in the range of 5–10 mg.

#### Differential scanning calorimetry (DSC)

Calorimetric measurements were performed using a DSC Q100 (TA Instruments). The samples were examined under nitrogen flow (50 mL/min) in an aluminum hermetic pan. Sample weights were in the range of 4–12 mg. Heating was performed at 5°C/min from 0°C to 200°C.

The denaturation temperature (T_D_) was determined as the peak value of the corresponding endothermic event. The denaturation enthalpy value was calculated with respect to the weight of air-dried gelatin.

### Human Articular Chondrocytes (hACs) harvesting and expansion

Human chondrocytes (hACs) were harvested from healthy femoral condyles and tibial plateau cartilage obtained from donors who underwent total knee replacement (TKR) prosthesis surgery. Specifically, tissues were harvested from 3 female donors (64, 71, and 82 years old), selected on the basis of well-defined inclusion criteria (unicompartmental OA, no previous knee surgery, no relevant comorbidities). Surgery waste was carefully washed with sterile PBS and placed in a sterile plate containing dissection medium (Diss-M: DMEM high glucose - Gibco USA, Penicillin/Streptomycin/Amphotericin 1% v/v - Gibco USA, Fetal Bovine Serum 10% v/v - Gibco USA). Then, the macroscopically undamaged cartilage was carefully removed using a sterile scalpel avoiding to harvest the underlying subchondral bone, and moved into a new sterile plate containing Diss-M, and minced in 1 cm^2^ pieces. Subsequently, minced cartilage was collected into a sterile 500 ml, conical tube and incubated with a 5 mg/ml solution of Collagenase A (Worthington, UK) in Diss-M at 37 °C under mechanical agitation (250 RPM) overnight. Finally, cells were recovered, filtered through a 40 µm cell strainer, counted, and plated at 7.0 X 10^3^ cell/cm^2^ in a tissue culture-treated T75 flask. Chondrocytes were allowed to grow until 80% confluency using chondro-FBS medium (cFBS-M: DMEM high glucose - Gibco USA, Penicillin/Streptomycin/Amphotericin 1% v/v - Gibco USA, Insulin/transferrin/selenium 1% v/v - Thermofisher USA, Dexamethasone 0.1 μM - Sigma-Aldrich USA, L-proline 40 μg/ml - Sigma-Aldrich USA, Fetal Bovine Serum 10% v/v-Gibco USA).

### GelMA-hACs, GelMA/GelOXA-hACs scaffold biofabrication

hACs were gently detached by adding 5 ml of TryPLE (Gibco USA) on T75 flask and maintaining at 37 °C for 10 min. Cells were recovered via centrifugation at 1200 rpm for 7 min (rotor radius = 247 mm), and pooled. The cellular pellet was resuspended in a 10% w/v solution of GelMA (Cellink, USA) in 0.25% w/v Lithium phenyl-2,4,6-trimethylbenzoyl-phosphinate (LAP) (Cellink, USA)/PBS solution, avoiding the formation of air bubbles during the whole process. For the GelMA/GelOXa-hACs scaffolds, 10% w/v of GelOXA was added to the cell suspension. Finally, the cell suspension was dispensed into a silicon mold, and a UV-mediated crosslinking was induced by irradiating scaffolds with an UV lamp equipped with led at 395 nm (LEPRO, USA) for 3 min. Hemisphere-shaped scaffolds were biofabricated (Ø = 1.5 cm, thickness ̴ 0.7 cm) (figure SI4), and 3 independent replicates were performed for each condition.

### Scaffolds culturing

Scaffolds were carefully removed from the silicone molds and moved in a not-tissue culture-treated 24-well plate (day -1, figure 10). Then, each scaffold was provided with 1 ml of cFBS-M, and incubated at 37 °C in an atmosphere containing 5% CO_2_ for 24 h, at which point (day 0, figure 10) the cFBS-M was switched into chondro complete medium (cCM: DMEM high glucose - Gibco USA, Penicillin/Streptomycin/Amphotericin 1% v/v - Gibco USA, Insulin/transferrin/selenium 1% v/v - Thermofisher USA, Dexamethasone 0.1 μM - Sigma-Aldrich USA, L-proline 40 μg/ml - Sigma-Aldrich USA, Ascorbic acid 50 µg/ml - Sigma-Aldrich USA, TGF-β3 10 ng/ml - Peprotech UK, without FBS).

### Viability assays

Cells or cell-laden scaffolds were washed twice in PBS to remove residual culture media, then stained using a LIVE/DEAD assay (Abcam, UK). Samples were stained using 4 μM calcein and 2 μM Ethd-1 (final concentration) in PBS for 30 min at 37°C in an atmosphere of 5% CO_2_. After staining, samples were rinsed twice in PBS to remove excess dye and imaged using an EVOS M5000 inverted epifluorescence microscope (Thermofisher, USA).

### Proliferation assay

Alamar blue assay (Thermofisher, USA) was used to evaluate the proliferation rate of chondrocytes. To build a calibration curve, hACs were seeded at decreasing density from 1.0 X 10^5^ to 9.0 X 10^3^ cell/cm^2^. Samples from 2D cell culture (n = 3) were plated at 1.5 X 10^5^ cell/cm^2^ into a tissue culture-treated 96-well plate in cFBS-M (Day -2). After 48 h (Day 0), the cFBS-M was switched into cCM for cells to be used in the calibration curve as well as for 2D culture test samples. All samples for the calibration curve were analyzed on day 0. For 2D cell culture, measurements were performed daily from day 0 to day 7 following the manufacturer’s instructions. Briefly, a 10% v/v dilution of Alamar blue in cCM was used to incubate cells for 3 h at 37 °C with 5% CO_2_, and measurements of fluorescence (λ_Exc_ = 560 nm, λ_Em_ = 590 nm) were carried out using a multiplate reader Spark 10M (Tecan, Switzerland).

### Histology

Cells cultured in 2D were fixed for 30 min at 4°C in 4% paraformaldehyde (Sigma, USA), washed twice with PBS, and stored in PBS for immunocytochemistry assays. Cellularized scaffolds were fixed overnight in 4% v/v paraformaldehyde (Sigma, USA) at 4°C under gentle agitation. Constructs were then dehydrated in an ascending ethanol series (25%, 50%, 70%, 95%, and 100%) before embedding in histology-grade paraffin (Leica biosystems, Germany) using a HistoCore Arcadia H embedding station (Leica biosystems, Germany). Then, embedded samples were sectioned at 5 μm thickness using a fully-motorized rotary microtome Leica RM2265 (Leica biosystems, Germany), collected on superfrost plus adhesion slides (Thermofisher, USA), and stored at room temperature until immunostaining.

### Immunocytochemistry

Immunostaining was performed on 24-well multiwell plates at days 0 and 7 for both 2D cell culture, and cell-laden scaffolds. Briefly, cells/scaffolds were washed twice with PBS, and treated with 70% v/v EtOH for 15 min at RT to fix cells. Suppression of nonspecific binding, and cells membrane permeabilization, were performed using a solution of 0.1 % w/v Triton X-100 (Sigma-aldrich, USA) and 1% w/v bovine serum albumin (BSA) (ITW, USA) in PBS for 45 min at RT. Samples were then washed twice and incubated with primary antibodies against type II collagen (ab34712, Abcam, UK), and type X collagen (ab49945, Abcam, UK), using a 1/50 v/v dilution overnight at 4°C. Finally, samples were washed three times with PBS, and incubated with fluorescent secondary antibody (Ab96919, and Ab98795 respectively, Abcam, UK) using a 1/50 dilution together with 4’,6-diamidino-2-phenylindole (DAPI) as nuclear staining. Samples were visualized using an epifluorescence microscope Evos M5000 (Thermofisher, USA).

### Histochemistry and immunohistochemistry

Scaffolds were fixed overnight in 4% w/v paraformaldehyde (Sigma, USA) at 4°C under gentle agitation, followed by dehydration in an ascending ethanol series (25%, 50%, 70%, 95%, and 100%). Then, a soaking in Xylene was performed prior the embedding in histology-grade paraffin (Leica biosystems, Germany) using a HistoCore Arcadia H embedding station (Leica biosystems, Germany). The embedded samples were sectioned at 5 μm thickness with a fully-motorized rotary microtome Leica RM2265 (Leica biosystems, Germany), collected on superfrost plus adhesion slides (Thermofisher, USA), and stored at room temperature in the dark.

For histochemical and immunohistochemical staining, the sections were de-paraffinized with Xylene (Sigma, USA), rehydrated by soaking in a descending ethanol series (100%, 95%, 70%, and 50%), and gently washed under deionized water for 2 min.

#### Alcian blue staining

Slides were soaked in an Alcian blue solution 1% w/v in acetic acid (pH = 2.5) (Sigma, USA) for 1h at RT, and the staining solution was washed away under deionized water. Then, slides were dehydrated in ascending ethanol series (25%, 50%, 70%, 95%, and 100%), mounted with organ/limonene mount medium, and stored at room temperature away from light.

#### Immunohistochemistry

For immunostaining, Ms-HIF1α (dilution 1:100, ab16066, Abcam, USA), Rb-COLI (dilution 1:42 v/v, ab233080, Abcam, USA), Rb-COLII (dilution 1:50 v/v, ab34712, Abcam, USA), and Ms-COLX (dilution 1:1000 v/v, ab49945, Abcam, USA) were used as primary unconjugated antibodies. Donkey anti-Mouse DL550 (dilution 1:50 v/v, Ab98795, Abcam, USA), and Donkey anti-Rabbit DL488 (dilution 1:50 v/v, Ab96919, Abcam, USA) were used as secondary conjugated antibodies for visualization. Propidium iodide (dilution 250 µg/ml, ab14083, Abcam, USA), and 4’,6- diamidino-2-phenylindole, dihydrochloride (DAPI) (Thermofisher, USA) were used as nuclear staining. Briefly, antigen retrieval was performed using an aqueous citrate buffer at pH 6.0 containing Sodium Citrate dihydrate 0.294% w/v (Sigma, USA), 5 µl Tween 20 (Sigma, USA), and deionized water.

Slides were then incubated within a StainTray slide staining system (Merck, Germany) with primary antibodies overnight at 4°C in the dark. Then, the primary antibody was washed away by rinsing three times in PBS, and samples were incubated with a mixture of secondary antibody and nuclear staining for 30 min at RT in the dark. Finally, slides were mounted with ProLong™ Glass Antifade Mountant (Thermofisher, USA) and stored in the dark at 4°C.

### Statistics

All analyses were performed using Prism 9.0 (GraphPad software, USA). The experimental results are expressed as means ± standard deviation (SD). Unpaired *t*-test with Welch’s correction, One-way ANOVA test followed by Tukey’s *post-hoc* test, or Friedman test followed by Dunn’s *post-hoc* tests were used, basing on preliminary analyses of data distribution. Differences were considered statistically significant if p < 0.05 (*p < 0.05, **p < 0.01, and ***p < 0.001).

## RESULTS

### OXA, and GelOXA derivatives synthesis, and chemical-physical characterization

#### High yields of OXA, and GelOXA compounds

Perfluoroheptylamidoxime **1** was treated with pentafluorobenzoyl chloride in the presence of pyridine as base. The reaction was performed in toluene, avoiding the use of benzene as a solvent, as previously reported^51^ (Figure 1A).

**Figure 1:**
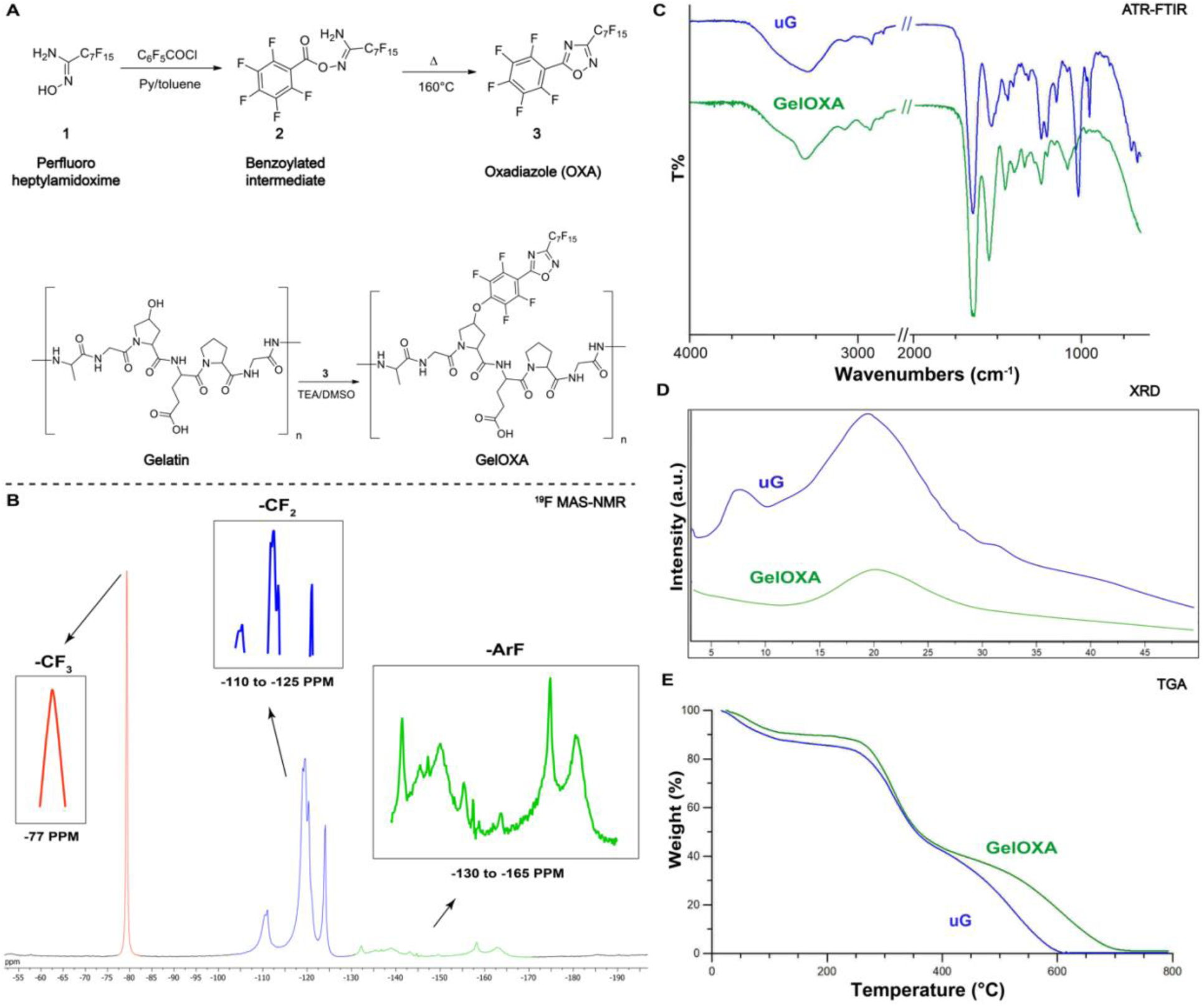
Synthesis of fluorinated 1,2,4-oxadiazole (OXA), and fluorinated gelatin-oxadiazole (GelOXA) derivatives (A). ^19^F Magic angle spinning (MAS) solid-state NMR (ssNMR) spectrum of GelOXA (B). Attenuated total reflectance Fourier transform infrared (ATR FT-IR) analysis of unmodified gelatin (uG) (blue) and GelOXA (green) (C). Wide-angle X-rays diffraction (XRD) patterns unmodified gelatin (uG) (blue) and functionalized gelatin (green) (D). Thermogravimetric analysis (TGA) curves of unmodified gelatin (uG) (blue) and functionalized gelatin (GelOXA) (green) (E).

The benzoylated compound **2** was obtained in excellent yield after simple filtration and reacted at high temperature under solvent-free conditions, to give oxadiazole **3** (OXA) in good yield without chromatographic separation (Figure 1A). The obtained oxadiazole **3** reacted with Tipe A gelatin in DMSO as the solvent and triethylamine (TEA) as the base (Figure 1B). Under these conditions, compound **3** can react with the nucleophilic moieties of gelatin, mainly -OH groups of hydroxyproline. Functionalized fluorinated gelatin (GelOXA) was obtained in excellent yield after precipitation in ethyl acetate. The solid GelOXA was simply recovered after centrifugation, washed, and centrifuged twice.

#### GelOXA characterization via ^19^F MAS-NMR, ATR FT-IR, XRD, and thermogravimetric analyses

Different techniques were employed to verify gelatin’s functionalization and assess GelOXA properties. ^19^F Magic angle spinning (MAS) solid-state NMR (ssNMR) is reported in Figure 1B, which clearly shows the presence of the fluorinated moieties of the oxadiazole compound. The -CF_3_ group (around -77 ppm), the -CF_2_-groups (in the region from -110 to -125 ppm), and the fluorine atoms linked to the aromatic ring (in the region from -130 to -165 ppm) are clearly visible. The infrared spectra recorded using an attenuated total reflectance Fourier transform infrared (ATR FT-IR) spectrophotometer of unmodified gelatin (uG), and of uG conjugated with fluorinated oxadiazole (GelOXA), are reported in Figure 1C.

The absorption bands characteristic of gelatin, associated with the vibrational modes of peptide bonds, are well recognizable at 1634, 1525, and 1237 cm^-1^ (blue line), and refer to referred as Amide I, Amide II, and Amide III, respectively. The intense band of Amide I, the band of Amide II, and the band of Amide III, are originating from the -C=O stretching vibration, -NH bending vibrations, and -CN stretching respectively^52^. Moreover, Amide A (N–H stretching) at 3290 cm^-1^, and Amide B (C-H stretching) bands at 3076 cm^-1^ are also visible. The spectrum of GelOXA (green line) is enriched by the presence of sharp absorption bands ranging from 900 to 1200 cm^-1^, characteristic of C-F bonds resulting from the perfluoroalkyl chain, and the perfluorophenyl ring of the conjugated groups. Furthermore, the broadening of the band centered at about 1525 cm^-1^ is likely due to the overlap of Amide II with the bands of the phenyl ring, thus confirming the functionalization^47^.

Figure 1D shows the wide-angle X-rays diffraction (XRD) patterns recorded for both uG and GelOXA. The pattern of uG (green line) includes two broad diffraction reflections: the first one, centered at about 8°/2 theta, is related to the diameter of the triple helix, while the second halo at about 20°/2 theta, corresponding to a periodicity of about 0.45 nm, is related to the distance between amino acidic residues along the helix^52^. In the pattern collected from GelOXA, the reflection centered at 8°/2theta disappeared, and the area of the reflection centered at about 20° clearly decreased, indicating the absence of an ordered structure.

These results are coherent with those from the calorimetric measurements, as the denaturation temperature of uG was set at 76°C, about 7 degrees higher than that recorded for GelOXA.

The thermal stability of either uG, and GelOXA, was evaluated by thermogravimetric analysis, as reported in Figure 1E. Gelatin is decomposed through multiple decomposition steps. The initial weight loss (up to 200°C) corresponds to the elimination of absorbed, and bounded water. The main degradation step, between 200°C and 450°C, is a complex process including protein chain breakage and peptide bond rupture, whereas at higher temperatures the degradation and combustion of organic residues occurs.

Both uG, and GelOXA, contained a similar amount of water (10% and 14% wt), but the onset of water loss occurred at 49°C and 33°C respectively, indicating a greater presence of unbounded water in the functionalized sample. Even the onset of the third process of GelOXA occurred at a lower temperature (487°C vs 538 °C), suggesting a lower thermal stability of the products of degradation.

### Our harvesting technique, and the use of cFBS-M, preserves hACs native phenotype and their viability after harvesting, and expansion under normoxic conditions

Our high-efficiency harvesting technique has been demonstrated to maintain hACs viability, as evidenced by their viability at 24 h (day 0) (figure SI1 A, B) after harvesting and after 7 days (figure SI1 C, D) of *in vitro* culturing in 2D. We evaluated cell viability using a Live/Dead assay. Although a slight reduction in viability was noticed at day 0, with a difference in the number of viable cells between the border and the center of the wells, the overall viability of cells increased over time up to day 7, with a corresponding reduction in the difference in viability between the border and the center of the well (figure SI1 E).

The minimal changes in cell density observed via brightfield microscopy, suggest that hACs do not exhibit significant proliferative activity throughout the experimental time frame. Cell density appeared to be slightly altered up to day 2, followed by a stabilization over time up to day 7 (figure 2A). These findings are consistent with results from the Alamar blue proliferation assay, which indicates a non-significant modification of the native non-proliferative phenotype of hACs (figure 2B).

**Figure 2:**
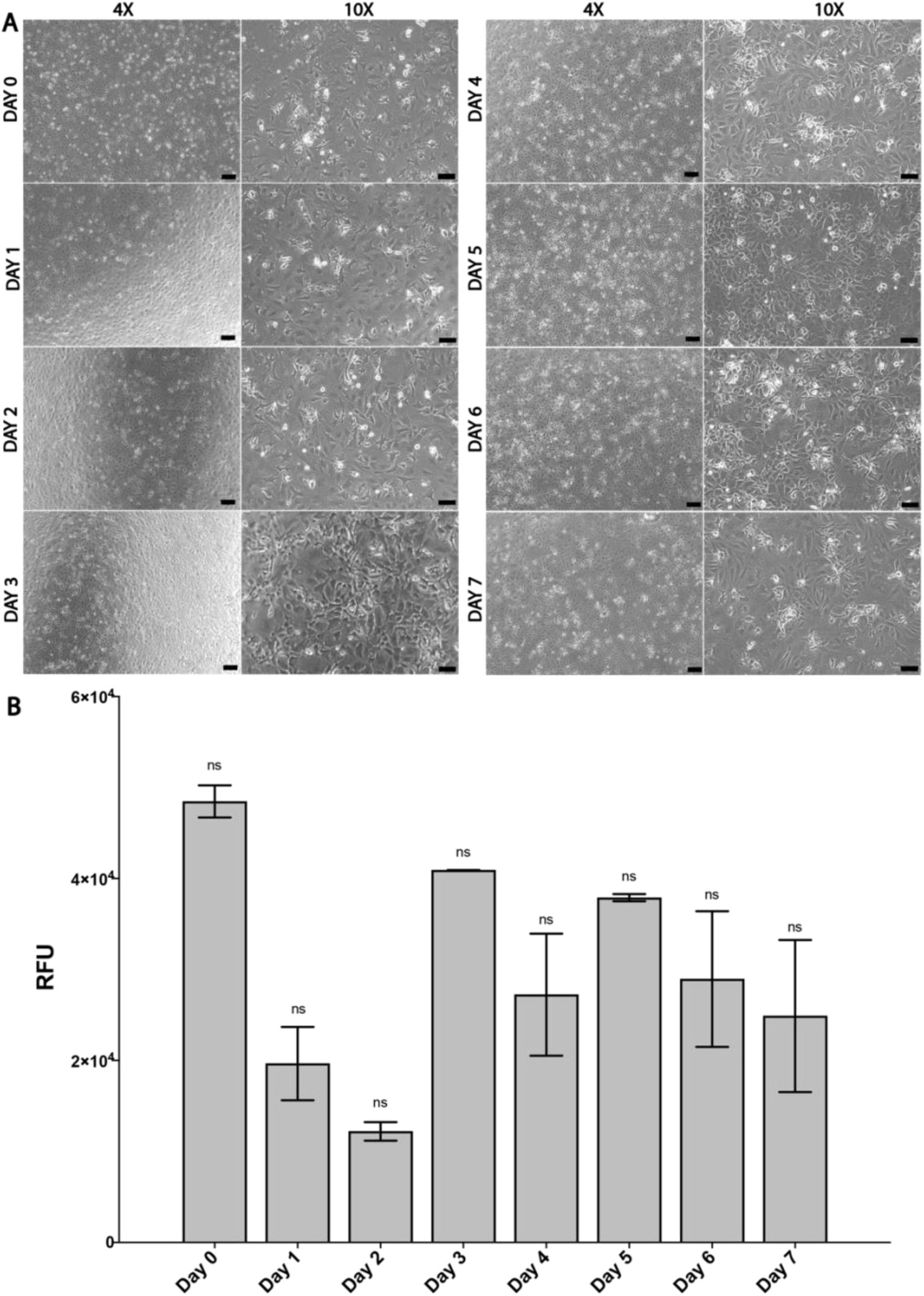
hACs cultured in a 24-well plate imaged daily in brightfield from day 0 to day 7 using a 4X (scale bar = 200 µm) and a 10X (scale bar = 100µm) objective (A). Proliferation assay for hACs during 2D culturing for 7 days quantified using the Alamar blue assay (B). Data are reported as mean ± SD, n = 3. Friedman test followed by Dunn’s *post-hoc* test, ns = non-significant.

The hACs showed an overall good capacity to produce GAGs, which are among the most representative ECM components of hyaline cartilage^53^. The GAGs were identified after staining with Alcian Blue, a group of water-soluble polyvalent basic dyes that can bind to sulfated and carboxylated mucopolysaccharides, and sialomucins resulting in a light to intense blue pigmentation. Notably, the slight but detectable blue staining observed in hACs immediately after harvesting on day 0 (figure 3 A, B), increased after 7 days of culturing in 2D (day 7) (figure 3 C, D).

**Figure 3:**
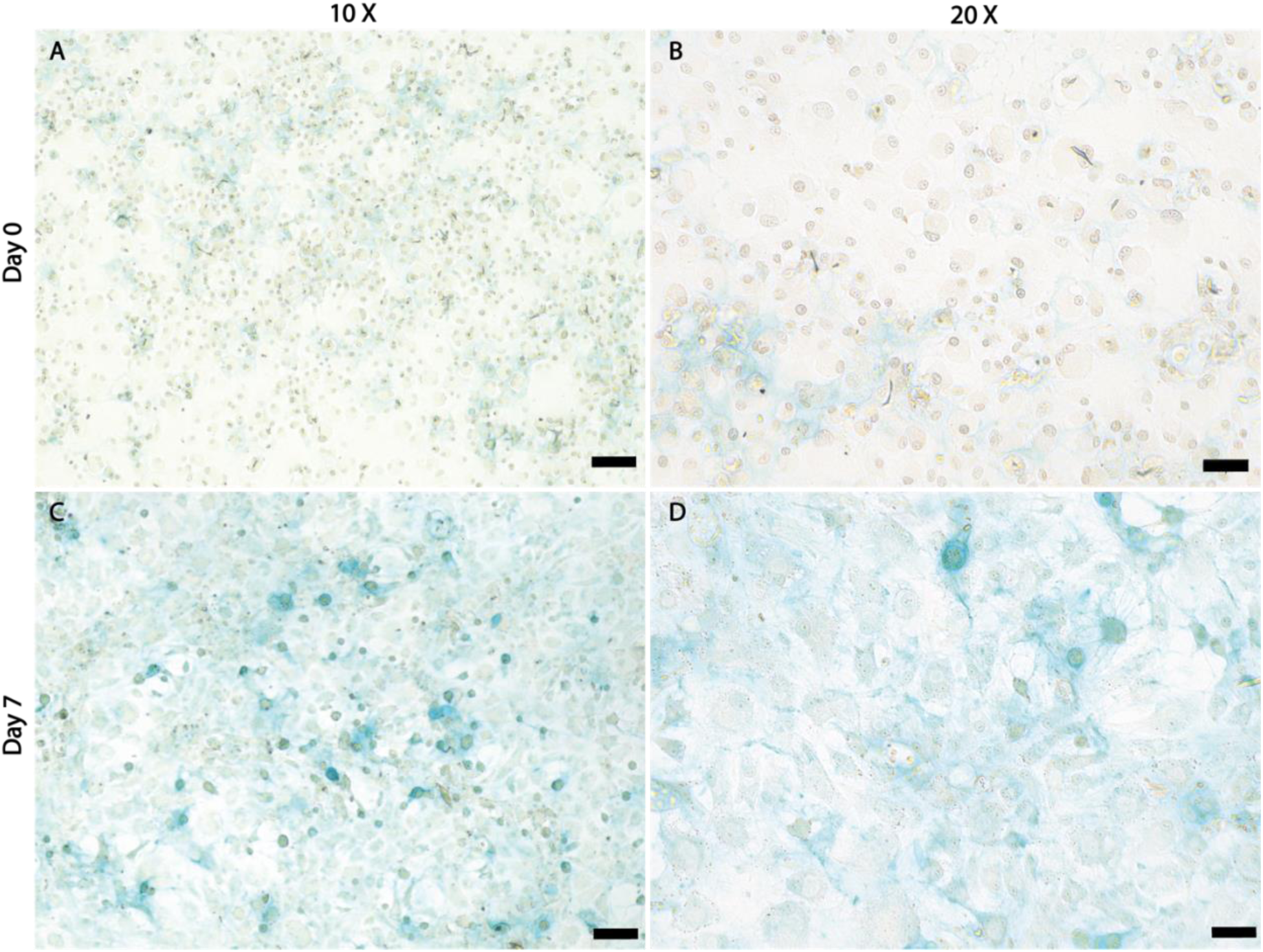
Alcian blue staining of hACs cultured in 2D at day 0 (A, B), and at day 7 (B, C) for GAGs quantification. Scale bar = 100 µm (A, C), and 50 µm (B, D).

The maintenance of the characteristic phenotype of hACs was confirmed by the presence of type II collagen, as demonstrated by immunocytochemical staining (figure 4 A, B). Notably, the amount of type II collagen significantly increased after 7 days of culturing (figure 4 E), with visible spread in the peri-cellular and extracellular space (figure 4 C, D). These findings strongly indicate a positive effect of our expanding protocol using c-FBS-M on the preservation of the native hACs phenotype, which produces extracellular matrix rich in type II collagen.

**Figure 4:**
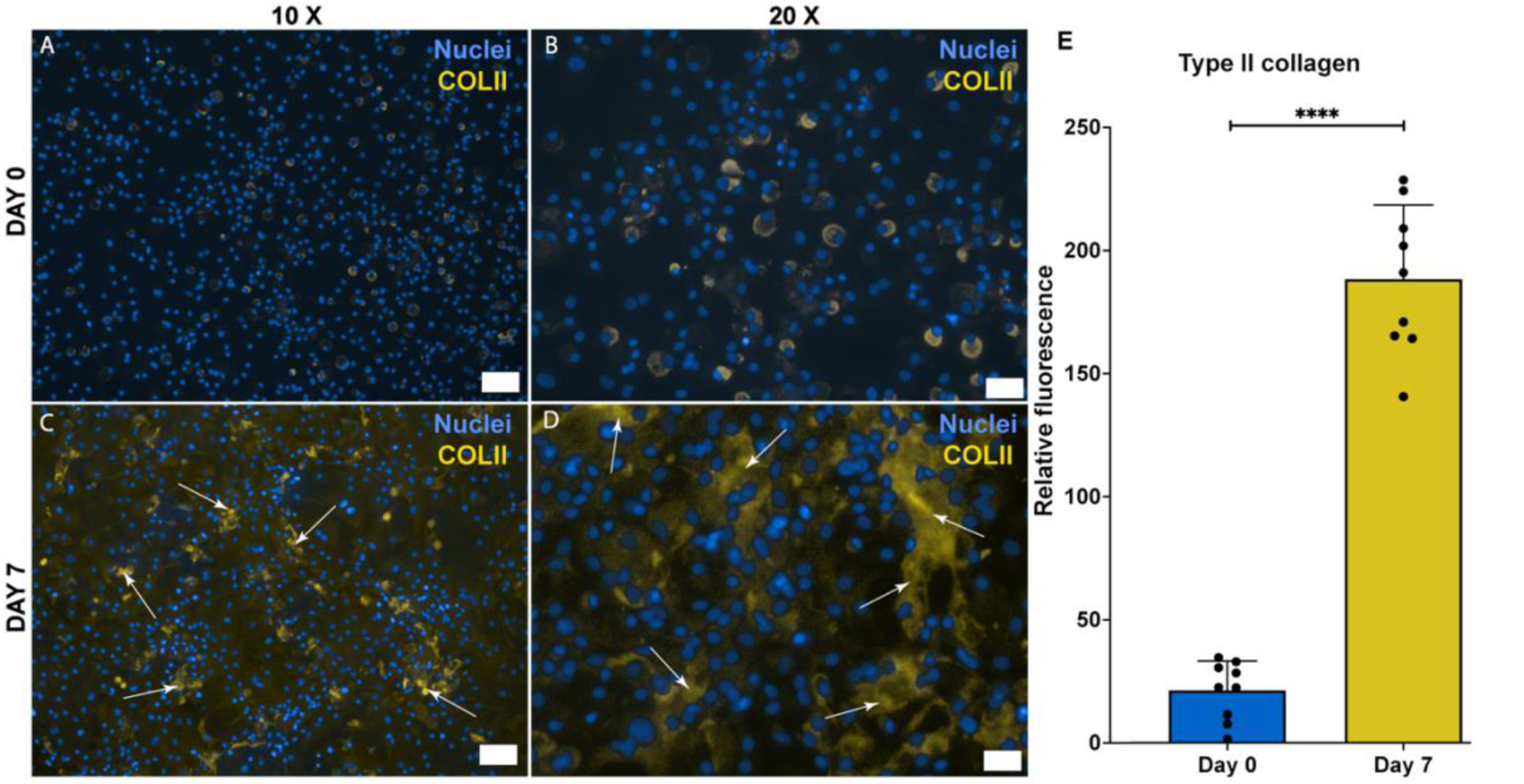
Type II collagen production from hACs cultured in 2D at day 0 (A, C), and at day 7 (B, D) imaged after immunohistochemistry. Yellow staining = type II collagen. Blue staining = nuclei. White arrows = peri-/extra-cellular spreading of type II collagen. Scale bar = 100 µm (A, C), and 50 µm (B, D). Quantification of type II collagen at day 0, and day 7 (E). Unpaired *t*-test with Welch’s correction, **** p < 0.0001. For the quantification, 9 images were randomly acquired over three independent replicates (n = 9).

### The GelMA-GelOXA scaffolds sustain cell viability in normoxic conditions

The viability of hACs in 3D culture was assessed using the same Live/Dead kit used for 2D culturing. The hACs exhibited robust viability immediately following scaffold biofabrication on day 0, as clearly indicated by the abundance of green spots visible in figure 5A. However, cells in the GelMa scaffold displayed reduced viability after 9 days of 3D culture, as evidenced by the increased number of red spots present after both hypoxic and normoxic culturing (figure 5B, C). Conversely, culture in the GelMA/GelOXA scaffold resulted in a markedly higher number of green spots compared to red ones (figure 5D), suggesting that the GelMA/GelOXA scaffold provides an environment better suited for hACs viability in 3D.

**Figure 5:**
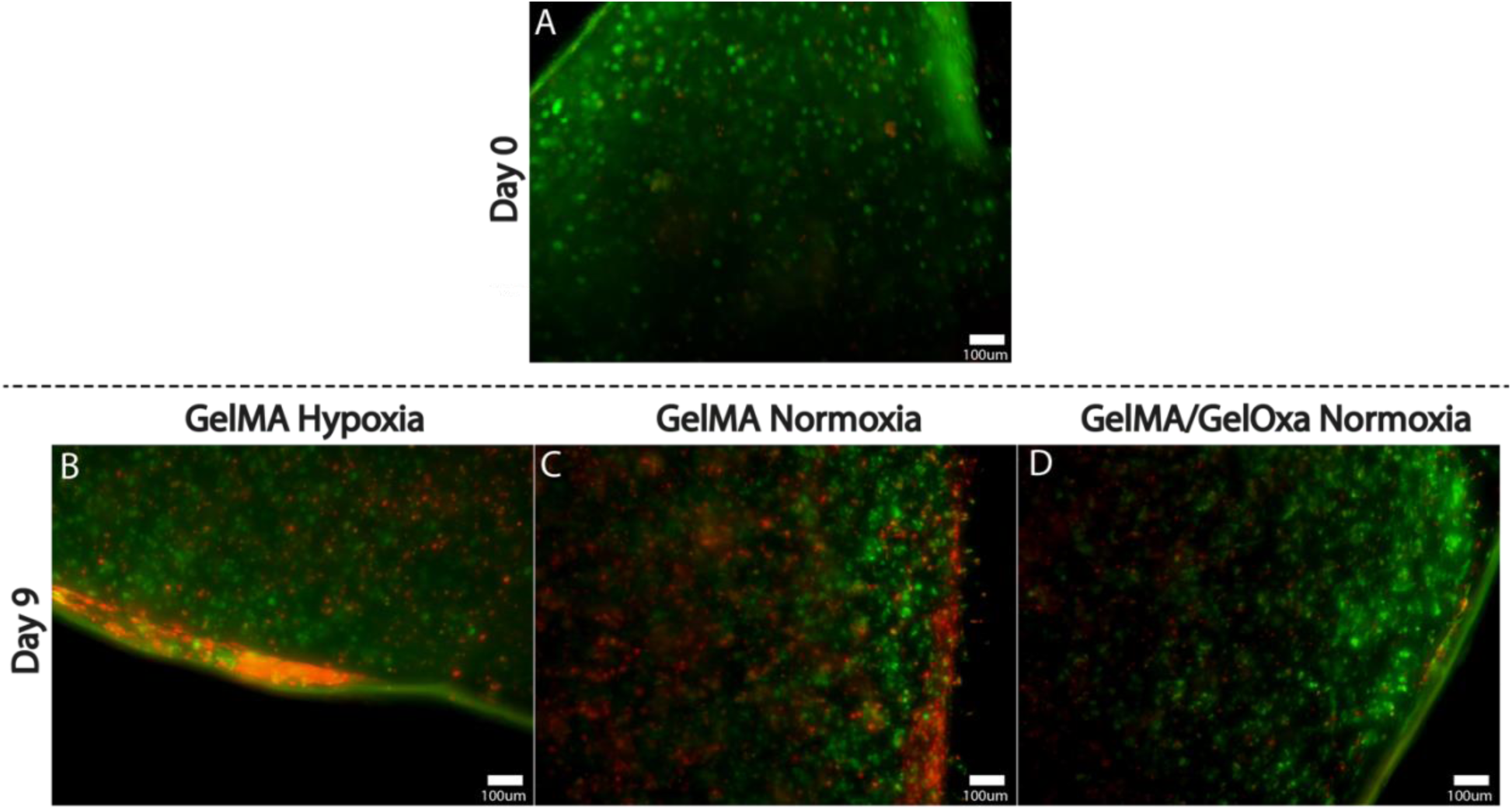
Live/Dead assay on hACs embedded in a 3D scaffold of GelMA at day 0 (A), and at day 9 after culturing under hypoxic (B), or normoxic conditions (C). Live/Dead assay on hACs embedded in a 3D scaffold of GelMA/GelOXA after 9 days of culture under normoxia. Scale bar = 100 µm.

### GelMA/GelOXA promotes the secretion of ECM characteristic of hyaline-like cartilage, via the activation of hypoxic-dependent pathways

Consistent with previous literature^54^, cells cultured in GelMA under hypoxic conditions showed greater production of type II collagen (figure 6A, D) than cells cultured in GelMA under normoxia (figure 6B, E). Remarkably, hACs cultured in our GelMA/GelOXA scaffold under normoxia displayed a remarkable capacity to produce type II collagen, with a significantly increased amount compared to GelMA in normoxia (negative control), as shown in figure 6C and 6F. Notably, there was no significant difference in type II collagen deposition between culture in GelMA under hypoxia (positive control) and our GelMA/GelOXA scaffold (figure 6G). This suggests that GelMA/GelOXA scaffolds can induce the production of a hyaline-like ECM rich in type II collagen even under normoxic conditions.

**Figure 6:**
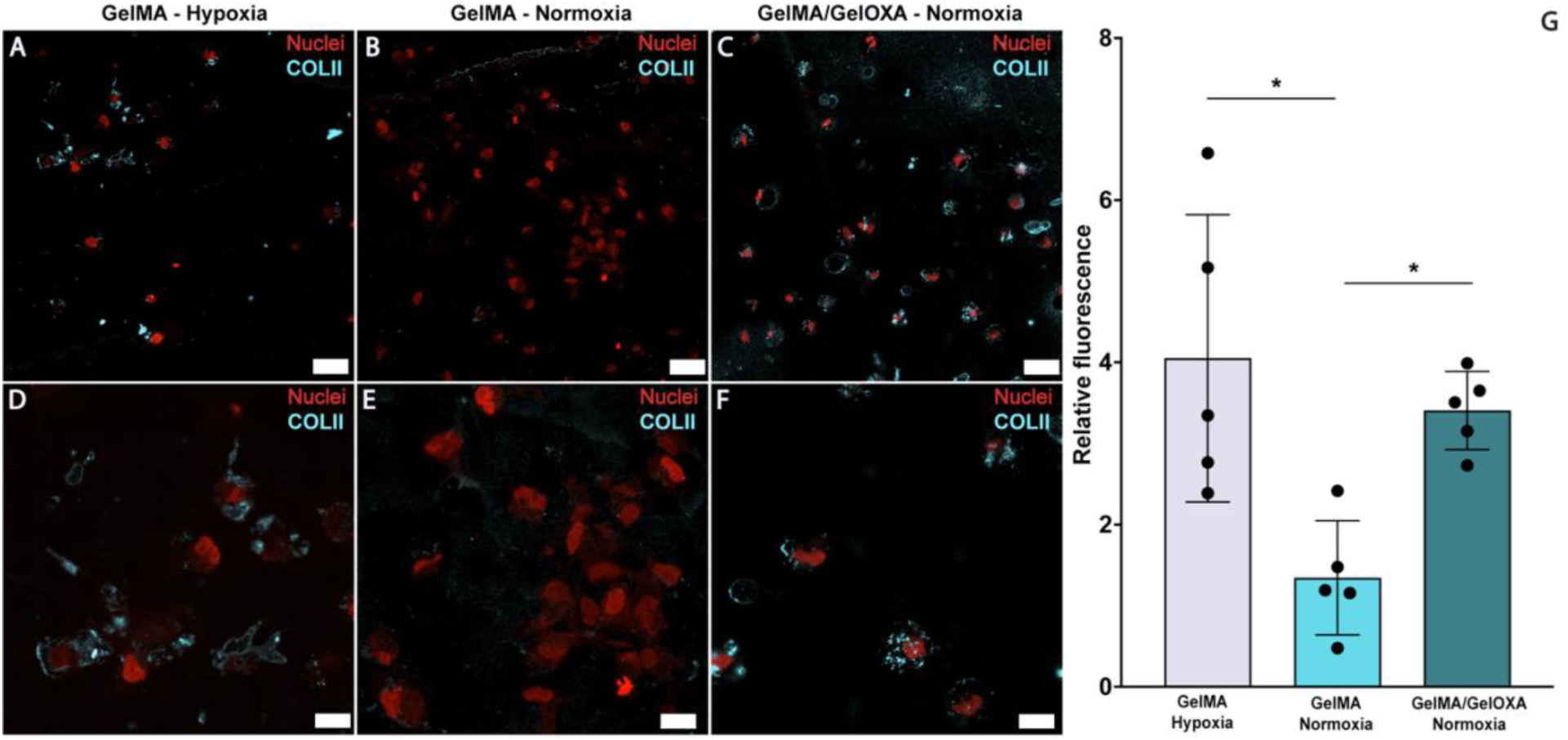
Images captured by confocal microscopy of hACs after 9 days of culture in GelMA scaffold in hypoxia (A, D), and normoxia (B, E). Immunohistochemistry on hACs after 9 days of culture within GelMA-GelOXA scaffold (C, F). Scale bar = 25 µm (A, B, and C). Light blue = type II collagen. Red = nuclei. Scale bar = 10 µm (D, E, and F). One-way ANOVA followed by Tukey’s *post hoc* test. For the quantification, 5 images were randomly acquired over three independent replicates (n = 5). *p < 0.05.

Conversely, results from the collagen X immunofluorescence showed that hACs embedded in GelMA cultured under both normoxic (figure 7B,E), and hypoxic (figure A,D) conditions, produced significantly lower amounts of type X collagen compared to hACs embedded in GelMA/GelOXA cultured under normoxic conditions (figure 7C, F). Interestingly, we observed the deposition of Collagen X, a marker selective for unhealth hypertrophic chondrocytes, in its characteristic perinuclear localization (SI3) in GelMA scaffolds cultured under normoxia (figure 7B, E, SI3).

**Figure 7:**
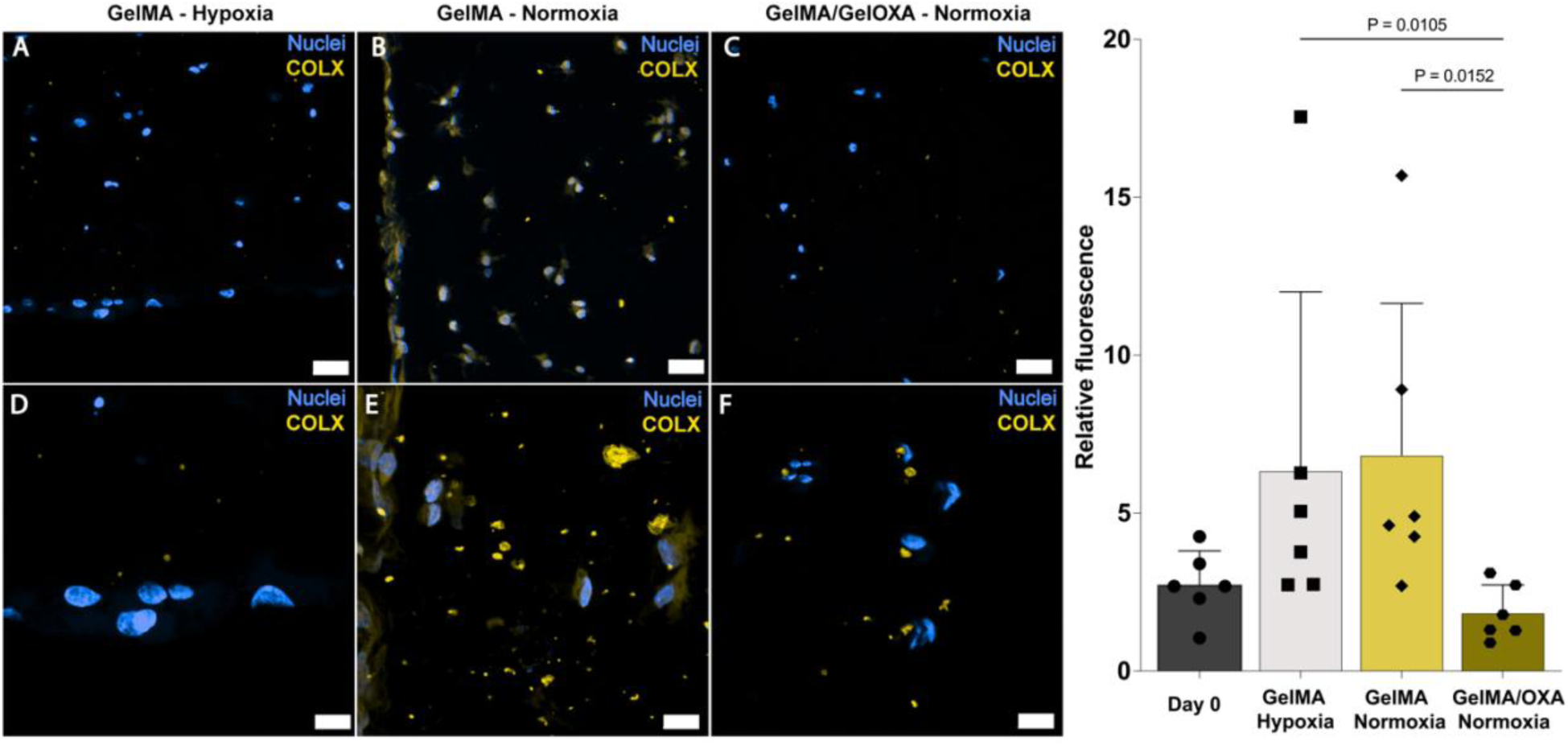
Images captured by confocal microscopy of hACs after 9 days of culture in a GelMA scaffold in hypoxic condition (A, D), and in normoxic condition (B, E). Immunohistochemistry on hACs after 9 days of culture in GelMA-GelOXA scaffold (C, F). Scale bar = 25 µm (A, B, and C). Yellow = type X collagen. Blue = nuclei. Scale bar = 10 µm (D, E, and F). Friedman test followed by Dunn’s *post-hoc* test. For the quantification, 6 images were randomly acquired over three independent replicates (n = 6). *p < 0.05.

The drifting toward a pre-hypertrophic phenotype of hACs embedded in GelMA scaffolds is also supported by the morphological modifications noticeable in images captured by SEM, showing a maintained native round-shaped morphology in hACs cultured in GelMA/GelOXA scaffolds (figure SI7).

Furthermore, the GelMA/GelOXA scaffold influenced the deposition of type I collagen, which is a hallmark of fibrotic cartilage. Specifically, hACs embedded in GelMA and cultured under hypoxic (figure 8A,D) and normoxic (figure 8B,E) conditions produced matrix containing similar amounts of type I collagen. Conversely, hACs cultured in the GelMA/GelOXA scaffold in normoxic conditions produced a significant lower amount of type I collagen compared to GelMA scaffold cultured in hypoxia (figure 8C,F).

**Figure 8:**
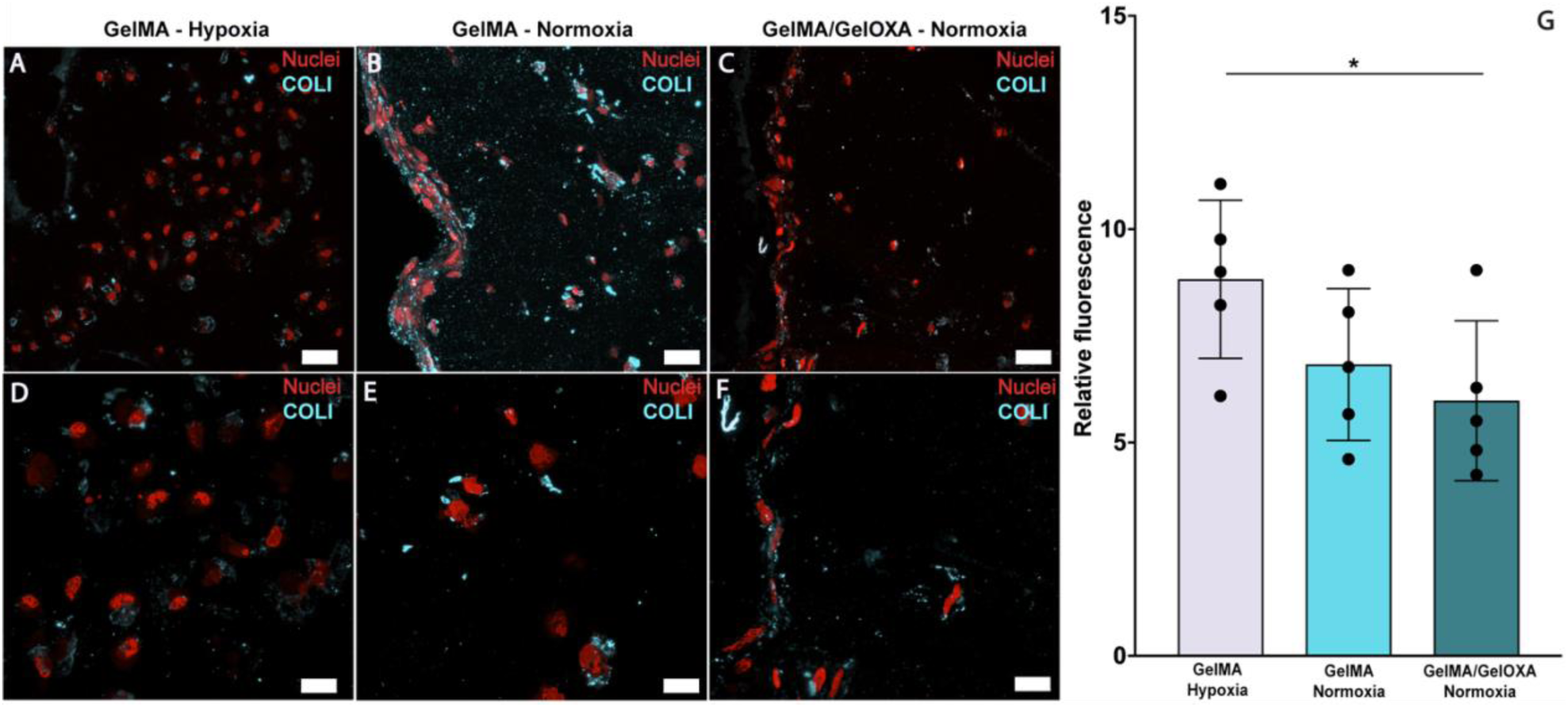
Images captured by confocal microscopy of hACs after 9 days of culture in GelMA scaffold in hypoxic condition (A, D), and in normoxic condition (B, E). Immunohistochemistry on hACs after 9 days of culture in GelMA-GelOXA scaffold (C, F). Scale bar = 25 µm (A, B, and C). Light blue = type I collagen. Red = nuclei. Scale bar = 10 µm (D, E, and F). One-way ANOVA test followed by Tukey’s *post-hoc* test. For the quantification, 5 images were randomly acquired over three independent replicates (n = 5). *p < 0.05.

The GelMA/GelOXA scaffold had a remarkable impact on the regulation of the hypoxia-driven HIF-1α pathway. In fact, hACs in the GelMA/GelOXA scaffold (figure 9C, F) induced the production of HIF-1α at levels similar to those produced by hACs cultured in GelMA under hypoxic conditions (figure 9A,D), and both presented significantly higher production (figure 9G) than the GelMA scaffold in normoxia (figure 9B,E). These findings strongly support our hypothesis that the GelMA/GelOXA scaffold is able to delivery hypoxic cues *in situ*.

**Figure 9:**
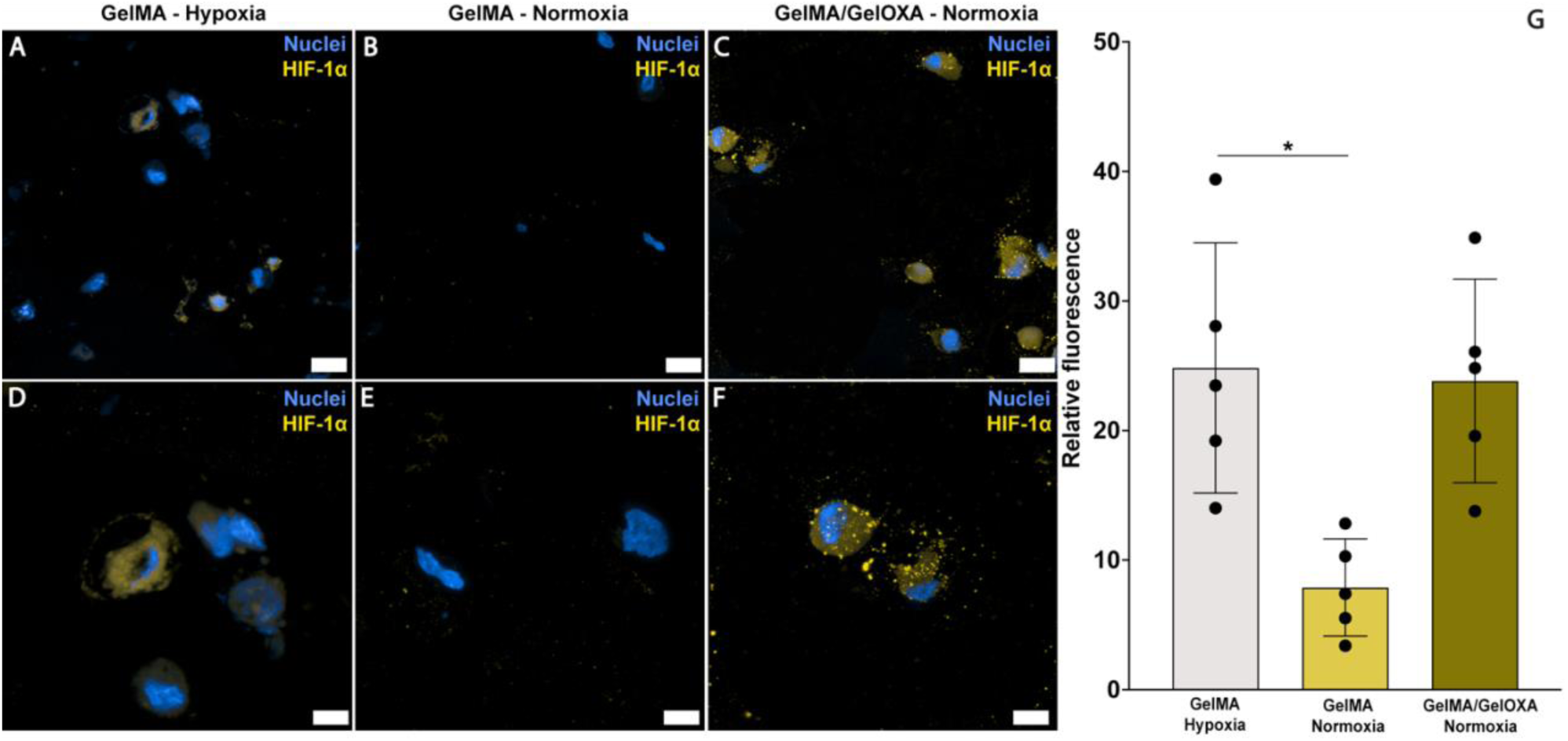
Images captured by confocal microscopy of hACs after 9 days of culture in GelMA scaffold in hypoxic condition (A, D), and in normoxic condition (B, E). Immunohistochemistry on hACs after 9 days of culture in GelMA-GelOXA scaffold (C, F). Scale bar = 25 µm (A, B, and C). Yellow = HIF-1α. Red = nuclei. Scale bar = 6 µm (D, E, and F). Friedman test followed by Dunn’s *post-hoc* test. For the quantification, 5 images were randomly acquired over three independent replicates (n = 5). *p < 0.05.

## DISCUSSION

Despite its simple cytological composition, hyaline cartilage (HC) is a dynamic tissue that responds and adapts to biochemical and mechanical stimuli to perform its physiological function, ensuring the correct functionality of the joints. Notwithstanding their slow metabolism, chondrocytes predominantly contribute to maintaining HC homeostasis and integrity. As a main undesired effect, their non-proliferative phenotype hinders joint self-healing, making damaged HC regeneration difficult without an external intervention.

For this reason, several efforts have been made to maximize the outcomes of RCTs, which are effective, but still cause undesired post-surgical modifications of the chondrocytes’ phenotype. This phenomenon usually occurs after the matrix implantation, triggering the chondrocytes de-differentiation^55^, ultimately leading to a decrease of the type II/type I, X collagen ratio^56^. This histologic arrangement is characteristic of fibrous/hypertrophic chondrocytes often observed in damaged HC affected by chondral defects^57,58^.

Several molecular mechanisms are reportedly involved in the etiology of these undesired changes in chondrocytes, with the cellular response to hypoxia being one of the most relevant. In fact, hypoxia is among the strongest promoters of type II collagen production in chondrocytes *in vivo* ^59^, an effect mediated by the s1α subunit of hypoxia-inducible factor (HIF-1α)^60–62^. In a normoxic environment, the oxygen-dependent enzyme prolyl hydroxylase domain (PHD) promotes the continuous proteasomal degradation of HIF-1α.

However, under hypoxic conditions, low oxygen concentrations inhibit PHD enzymes, ultimately leading to the stabilization of HIF-1α. The stabilized HIF-1α protein then translocates to the nucleus, where it forms a heterodimer with HIF-1β^63^, activating pathways critical for chondrocyte to adapt to the physiologically low-oxygen environment of the articular cartilage, supporting cell survival and maintaining overall cartilage homeostasis^64^.

Building on these knowledges, scientists have developed advanced scaffolds exploiting the positive effect of hypoxia in preserving the native chondrocytes phenotype in joints. The overall goal was to create a hypoxic environment within the scaffold, to induce the production of type II collagen from adult chondrocytes, ultimately producing a pro-regenerative effect.

Desferoxamine (DFO)^65,66^, or inorganic ions^67^, have been used to decrease the oxygen level within these scaffolds. However, several drawbacks have reportedly limited the application of these approaches, including systemic toxicity^68^ and the induction of DNA damages through oxidative-stress^69^.

Our work is conceived to overcome the limitations of these technologies, aiming to bridge the gap between beneficial and detrimental effects. Specifically, in this study we report the development of a novel bioactive scaffold capable of delivering pro-regenerative hypoxic cues to embedded hACs. Such a design is intended to re-establish the native hypoxic environment of the hyaline cartilage which is lost in joints affected by CDs. The main goal is to preserve the native chondrocytes phenotype, ultimately promoting the repair of chondral lesions *in situ*. The scaffold relies on a core composed of methacryloyl gelatin (GelMA), a gelatin derivative suitable for UV-mediated crosslinking reaction that increases the native structural stability of gelatin. The GelMA core is enriched with hypoxia-inducing seeds (GelOXA) composed of unmodified gelatin (uG) functionalized with a fluorinated oxadiazole (OXA), a class of compounds characterized by high oxygen affinity.

Gelatin is a collagen-derived biocompatible polymer^70^ that exposes different active moieties useful for chemical functionalization with OXA, whose oxygen affinity is due to the presence of fluorine atoms directly or indirectly linked to a nitrogen-based heterocyclic core. Overall, such a chemical-structural architecture perfectly aligns with our highly-demanding experimental need to locally modulate the oxygen concentration.

We exploited the fluorinating properties of the OXA (figure 1A), whose long perfluoroalkyl chain (C_7_F_15_) linked to the 1,2,4-oxadiazolyc core allows easy fluorination via nucleophilic substitutions^71^.

Interestingly, previous studies have demonstrated the feasibility of functionalizing biocompatible polymer backbones using OXA derivatives^48,72^, with no reported influence on the overall biocompatibility of the polymers. Notably, these classes of OXA-functionalized polymers acquire an oxygen affinity that is closely related to the oxadiazole concentration/unit repetition (UR) ratio^73^.

Basing on these preliminary data, we designed a mild-conditions synthesis for GelOXA that afforded high-yield gelatin functionalization, as well as an easy recovery of the final product (figure 1A). After a chemical characterization via ^19^F MAS-NMR, and FT-IR (figure 1B, C) confirming the functionalization, we moved forward to a physical characterization of GelOXA starting from the study of its thermal stability. The latter was comparable to thermal stability we observed in uG, although GelOXA exhibited a less ordered structure (SI 2).

We attribute this behavior to an altered renaturation equilibrium of the uG due to the introduction of OXA in the gelatin backbone. In fact, it is well known that gelatin is a random coil protein derived from the partial hydrolysis of collagen, whose triple-helix structure is naturally stabilized by interchain hydrogen bonds, and Van der Waal interactions. Such a tertiary structure of the collagen is broken down during the hydrolysis^74^, and may be reestablished in gelatin during a spontaneous partial renaturation^75^.

In GelOXA seeds, the presence of OXA might hampers the renaturation mechanism, ultimately stabilizing the denatured isoform of uG. To test our hypothesis, we performed additional analyses using wide-angle X-rays diffraction (XRD), and differential scanning calorimetry (DSC). These techniques provide valuable insights into the structural organization of the material. Specifically, the diffraction pattern of renatured gelatin typically exhibits a broad reflection centered around 8°/2 theta, and is known to be related to the amount of triple-helical structures^74^.

Consistent with our hypothesis, XRD analyses revealed the absence of such characteristic renaturation peak in GelOXA seeds compared to uG (figure 1D).

This finding suggests for a disruption of the hierarchical triple-helical structure within uG upon OXA functionalization, potentially due to the formation of less organized supramolecular assemblies. DSC analyses further validated this concept (SI 2), as we observed a decrease of the denaturation temperature of GelOXA compared with that of uG. This evidence indicates a destabilization of the triple-helical conformation and a stabilization of the unfolded random coil conformation in uG induced by OXA moieties. This can be attributed to the steric hindrance effects of the OXA groups, which might hinder the re-establishment of hydrogen bonds and the Van der Waal interactions crucial for the triple helix formation. The results from TGA (figure 1E) align with the proposed mechanism, as the decrease in bound water content observed in GelOXA suggests a less hydrated, and potentially less renaturable, protein network.

After successfully confirming the uG functionalization with OXA, we proceeded with the scaffold biofabrication. The scaffold design relies on GelOXA seeds as the active hypoxia-inducing component, dispersed within a GelMA matrix, which provided the main structural support. We cellularized the scaffold with human articular chondrocytes (hACs) harvested, and cultured, using our customized protocol. Notably, we accurately conceived this procedure to prevent the hACs de-differentiation and the related native phenotype loss (figure 2). Precisely, we used two different culture media, chondro-FBS medium (cFBS-M), and chondro complete medium (cCM), to harvest/expand, and culture hACs respectively, switching strategically between them to minimize the well-known detrimental effects of FBS on the hACs phenotype^76^.

We performed either hACs isolation, and expansion using cFBS-M to reduce the cellular stress, support the cells viability, and to induce a reversible and transitory proliferative phenotype. Remarkably, the hACs expansion never exceeded three passages (P3) to minimize any undesired effect on the hACs native-phenotype.

At P3 we switched the cFBS-M in favor of cCM, and maintained cells in this medium for up to 7 days to promote the transition from a proliferative to a non-proliferative hACs phenotype. We observed a high cell viability at day 0 followed by a slightly increase by day 7 (figure S1). Interestingly, we did not notice neither changes in morphology (figure 2A), nor increasing in cell proliferation (figure 2B). Moreover, we detected a non-significant decrease in metabolic activity between days 1 and 2, which we attribute to the hACs response to the serum-free conditions. Overall, these evidences support the efficacy of our harvesting and culturing approach in preserving the native non-proliferative cytotype of hACs without affecting their viability.

To delve deeper into the secretive activity of hACs, we studied and characterized the newly produced ECM. We first implemented an analysis based on Alcian blue, a staining selective for the positively charged GAGs that are key components of the hyaline cartilage proteoglycans. Interestingly, we spotted a weak extracellular blue pigmentation at day 0 (figure 3A, B) that markedly increased after 7 days of 2D culture (figure 3C, D).

To further characterize the ECM, we performed a quali-quantitative analysis based on immunostaining selective for type II collagen, which is abundantly produced by healthy hyaline cartilage^77^. Strikingly, we noticed that hACs were able to produce a high amount of type II collagen, which significantly increased after 7 days compared with day 0 (figure 4E). In addition, the immunostaining revealed a marked shift of type II collagen localization, from predominantly intracellular on day 0 (figure 4A, B) to prominent extracellular on day 7 (figure 4C, D), confirming the active secretive activity of hACs. These findings strongly suggest that our procedure effectively preserved the chondral phenotype of hACs during harvesting and expansion in 2D cultures.

These encouraging results were fundamental to proceed with the 3D scaffold biofabrication step, as we isolated and expanded hACs using the same protocol before embedding them within scaffolds (biofabrication). Precisely, we switched media from cFBS-M to cCM immediately after the biofabrication (day -1) (figure 10) to allow cells to recover for 24h, and cultured up to day 9 replacing media every 48h.

**Figure 10:**
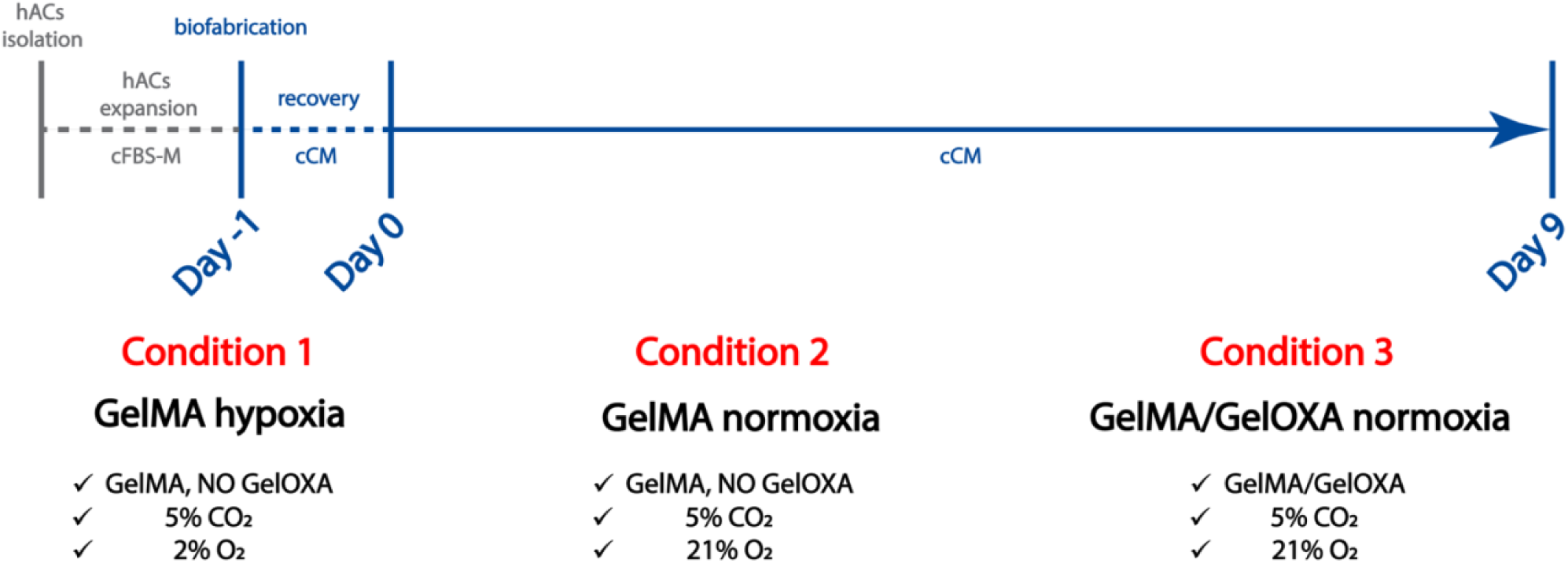
Timeline of the experimental design. Isolation and expansion were performed using a serum-enriched culture medium (cFBS-M). The cFBS-M was switched to a serum-free chondrospecific culture medium (cCM) after biofabrication at day -1. The experiment started after 24 hours of recovery time.

We thoroughly set up different experimental conditions to evaluate the response of hACs to a 3D culture within our hypoxia-inducing scaffolds. In condition 1 (figure 10 - GelMA hypoxia), we embedded hACs in a scaffold solely composed of GelMA, and cultured the system under hypoxic conditions in a hypoxic chamber. We conceived this condition as a positive control toward pre-established hypoxic cues. Conversely, in Condition 2, we included hACs in the same scaffold, but we cultured the system under normoxic conditions (figure 10 - GelMA normoxia) to create a comparative negative control. Condition 3 aimed to assess the capability of our GelMA/GelOXA scaffold to induce a pro-chondrogenic hypoxic-like environment. With this in mind, we embedded hACs in a GelMA/GelOXA scaffold and cultured the system under normoxic conditions (fig. 10 - GelMA/GelOXA normoxia).

We were impressed to observe a slight decrease of cell viability between day 0 and day 9 in GelMA cultured either in normoxic and hypoxic environment (condition 1, 2), in contrast to a dissimilar overall higher viability we have noticed in GelMA/GelOXA scaffolds (condition 3) (figures 5). Strikingly, although hypoxia has already been reported to increase chondrocytes viability^78^, here we demonstrate that our GelMA/GelOXA scaffold is more effective in sustaining hACs than a standardized hypoxic environment.

We hypothesize this might be the result of a localized hypoxia delivered by GelOXA seeds dispersed within the matrix, which may ultimately mimic the minimal O_2_ flow admitted by the synovial fluid circulation in the native cartilage *in vivo* ^79^ (dynamic hypoxia).

Moreover, the viability enhancement of hACs embedded within the GelMA/GelOXA scaffolds is coherent with the cells’ secretive activity, which is compatible with a hyaline-like phenotype. After 9 days, hACs embedded in the GelMA/GelOXA scaffolds cultured in normoxia (condition 3) exhibited a relevant type II collagen production, which is comparable to that of hACs embedded in gelMA cultured in hypoxia (condition 1) (fig. 6G). Notably, when cells were cultured in GelMA in normoxia (condition 2), type II collagen production was significantly lower than that in the GelMA/GelOXA scaffold under the same conditions (condition 3) (fig. 6G).

We observed an opposite trend in the secretion of either type X, and type I collagen, markers specific for hypertrophic, and fibrocartilaginous chondrocyte phenotypes respectively.

Specifically, type X Collagen is reportedly produced by terminally differentiated articular chondrocytes in diseased joints, and exhibits a characteristic peri-cellular localization *in vivo* ^80,81^. Interestingly, we found a type X collagen production with its characteristic perinuclear localization, (SI 3) from hACs embedded in the GelMA/GelOXA scaffold (condition 3), which was not significantly higher than type X collagen produced by hACs cultured in the GelMA scaffold under induced hypoxic conditions (condition 1). Remarkably, hACs produced a significantly higher amount of type X collagen when embedded in the GelMA scaffold (condition 2) cultured in same conditions used for the GelMA/GelOXA scaffold (figure 7G). We observed a similar secretion pattern for type I collagen, as it was significantly lower in the ECM of GelMA/GelOXA scaffolds cultured in normoxia compared to GelMA scaffolds cultured under both hypoxic and normoxic conditions (figure 8G).

Overall, these results indicate that the GelMA/GelOXA scaffolds support the maintenance of a native-like phenotype of hACs, as well as the deposition of an ECM rich in type II collagen, with not significant levels of type I/X collagen. These data, contextualized within our experimental conditions, strongly suggest the involvement of hypoxia-related mechanisms in GelMA/GelOXA scaffolds, and perfectly align with findings already observed *in vivo*^59^.

To further investigate this aspect, we analyzed the production of HIF-1α, a pro-regenerative factor in articular cartilage^82^ whose production is tightly correlated with hypoxic environments^64^. Notably, we found that hACs embedded in the GelMA/GelOXA scaffold cultured in normoxia (condition 3) produced an amount of HIF-1α comparable to that produced from hACs embedded in GelMA cultured under hypoxia (condition 1). Conversely, hACs embedded in GelMA cultured in normoxia (condition 2) produced a significantly lower amount of HIF-1α (figure 9G), thus testing our hypothesis regarding the hypoxia-inducing properties of the GelMA/GelOXA scaffold cultured under normoxic conditions.

## CONCLUSION

This study describes a significant step forward in the adoption of TE for cartilage regeneration using an innovative approach based on a scaffold capable to fine tuning hypoxia *in situ*.

Such scaffold exhibited the advantage of creating a controlled, and localized, environment that mimics the native oxygen levels of healthy articular cartilage. In addition, the GelMA/GelOXA scaffold we presented here showed to overcomes the limitations described for other hypoxia-inducing materials, which induce only transient hypoxia, and a short-lived stabilization of HIF-1α^83^.

Conversely, we demonstrated the efficacy of the GelMA/GelOXA scaffold in inducing, and maintaining HIF-1α up to 9 days. Such strong result is enforced by the evidence of increased production of type II collagen coupled with reduced expression of undesirable markers of hyaline cartilage (type I/X collagen). Strikingly, the scaffold promoted the deposition of healthy hyaline-like extracellular matrix even under normoxic conditions, suggesting its potential to deliver pro-regenerative cues directly at the injury site.

We plan to further improve this study by performing direct oxygen level measurements within the scaffold. Such an implementation will clarify the mechanism underlying the exciting results discussed so far.

Overall, the GelMA/GelOXA scaffold emerges as a promising strategy for cartilage regeneration for treating chondral defects. We believe that our approach can maximize the benefits of hypoxia in a controlled and targeted manner, holding the great promise for future successful cartilage regeneration.

## ACKNOWLEDGMENTS

This research was supported by the ON Foundation, starting-grant n°22-006. We thank you the Ri.MED Foundation for the partial contribution to this study. We are grateful to our colleagues Prof. Francesco Lopresti, and Prof. Vincenzo La Carrubba at the University of Palermo who provided a precious support that greatly assisted us for the SEM characterization.

## ETHICS STATEMENT

This study adheres to ethical principles and guidelines for research involving the use of primary human cells. Cells and tissues harvesting, as well as patients’ sensible data collecting, and storage, were performed according to a protocol approved by the ethical Institutional Research Review Board (IRRB) of the “Buccheri La Ferla” hospital.

