## Supplementary data for "Biofabrication of an *in situ* hypoxia-delivery scaffold for cartilage regeneration"

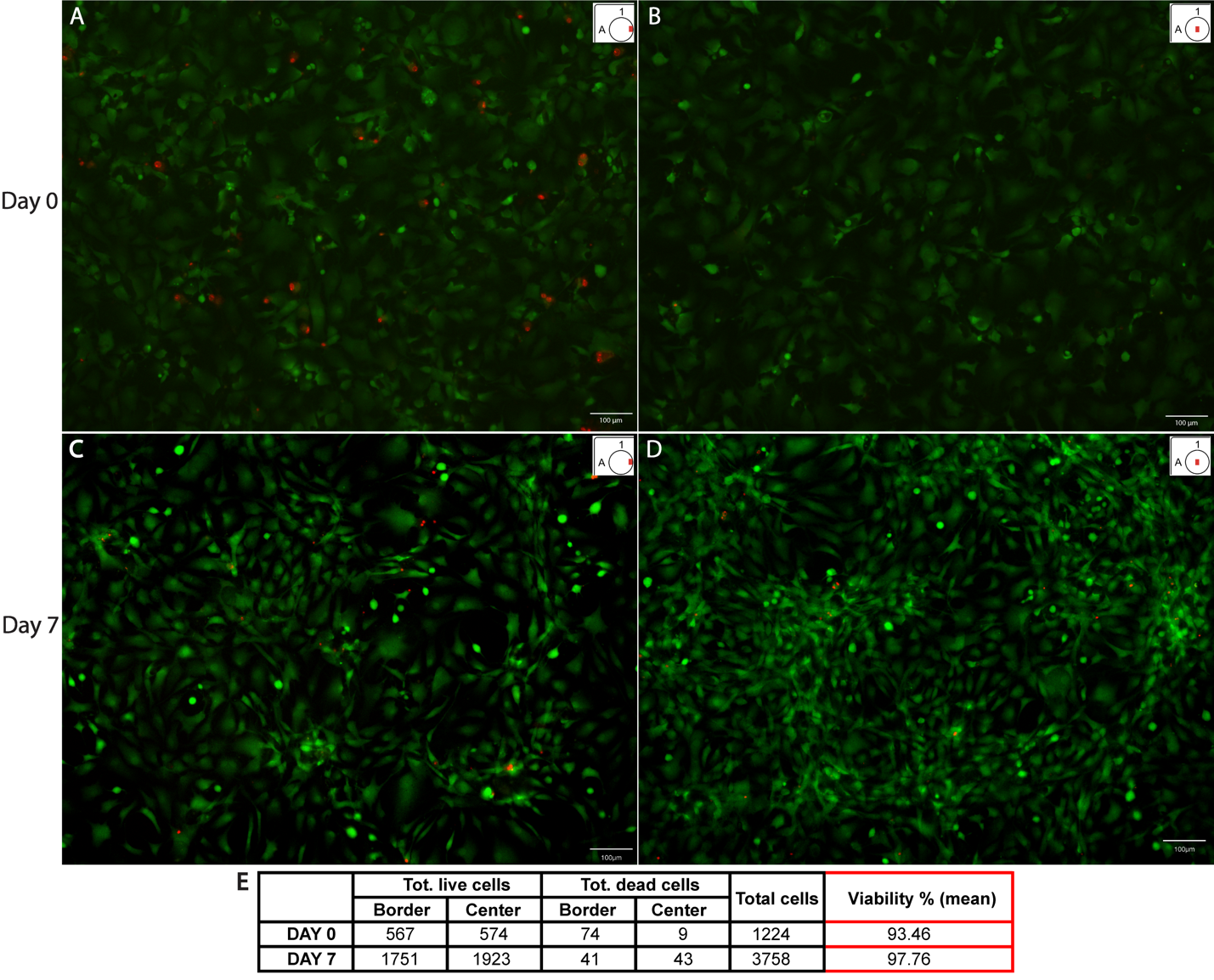

**SI 1:** Live/dead assay on hACs cultured within a tissue culture treated 24 well plate 24h (A, B) and 7 days (C, D) after the harvesting. Viable and dead cells are stained in green and red respectively. Scale bar = 100 µm.

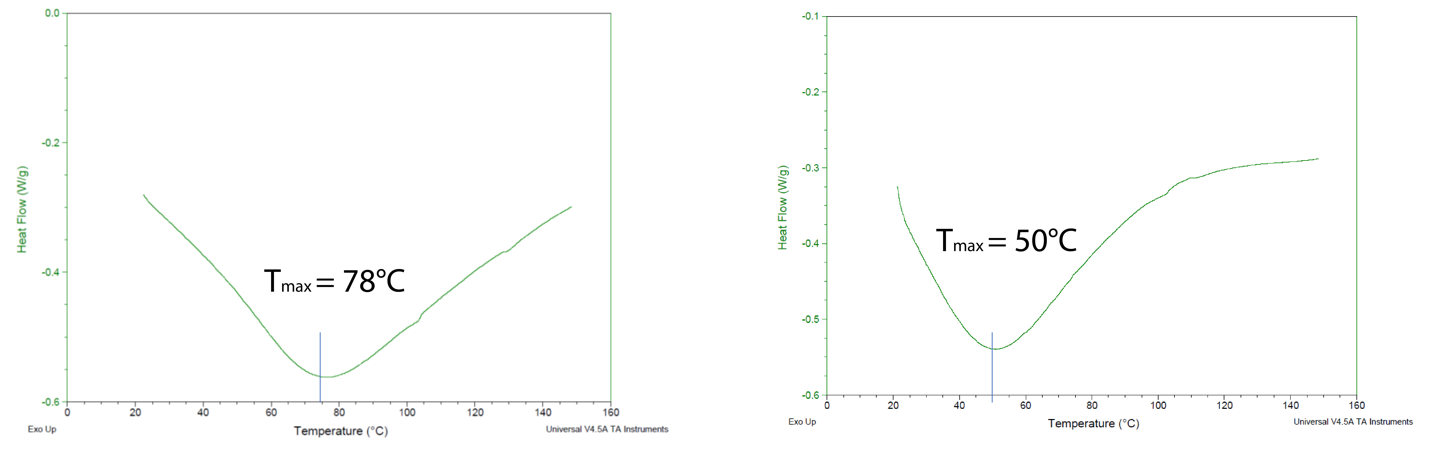

**SI 2:** Differential Scanning Calorimetry (DSC) analysis of uG (left), and GelOXA (right)

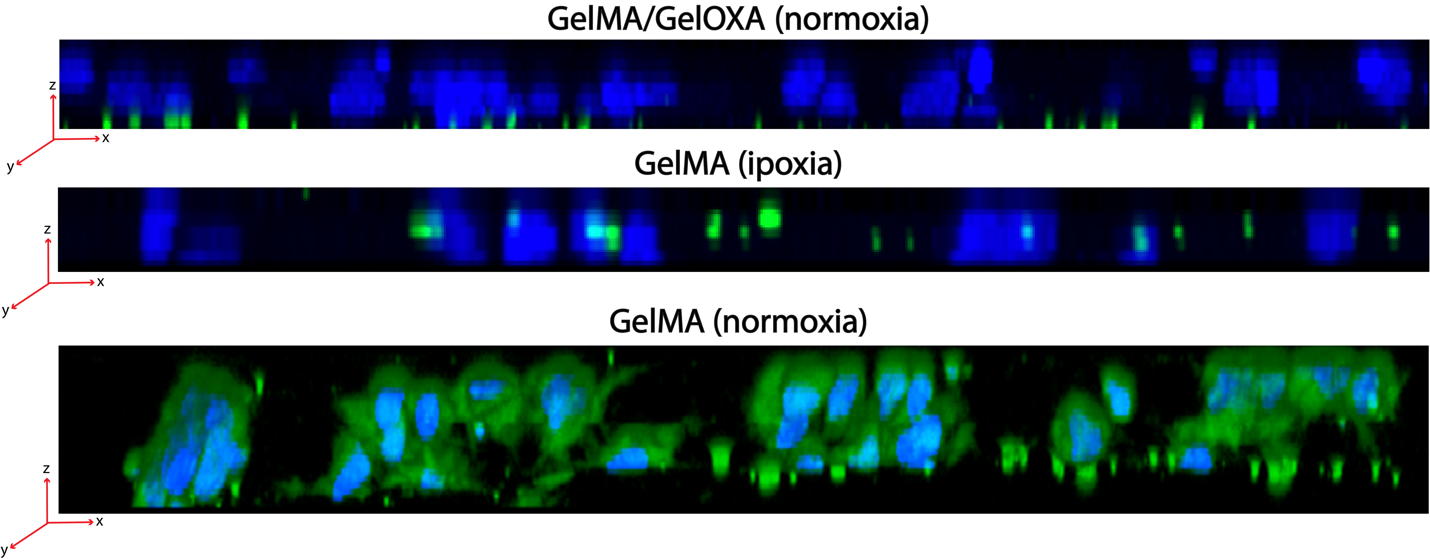

**SI 3:** 3D reconstruction of 2D planes obtained from Z-stack images acquired by confocal microscopy of type X collagen tissue distribution. Blue = nuclei, Green = type X collagen.

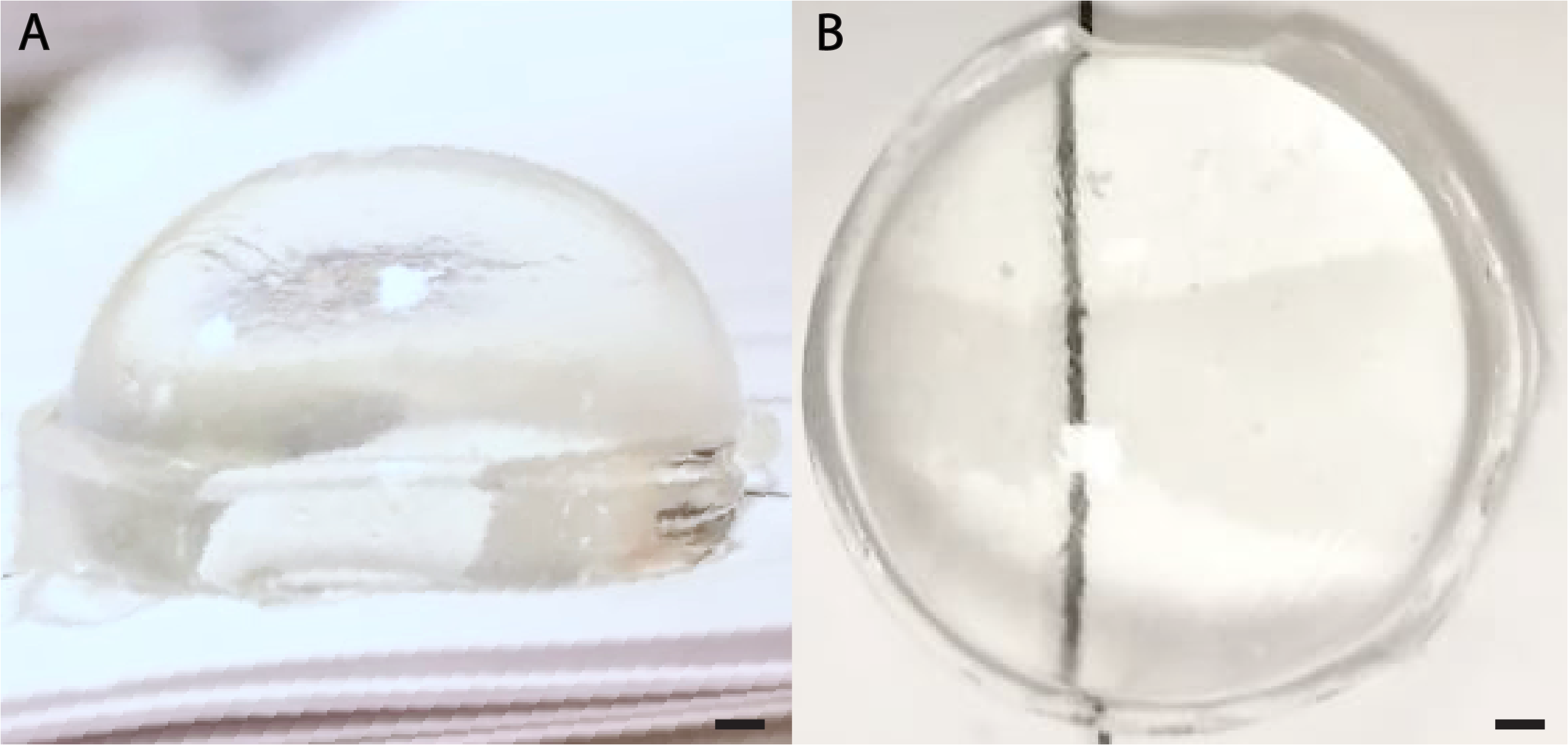

**SI 4:** Lateral (A), and top (B) view of an acellular GelMA scaffold immediately after the biofabrication. Ø = 1.5 cm, thickness ̴ 0.7 cm. Scale bar = 1 mm.

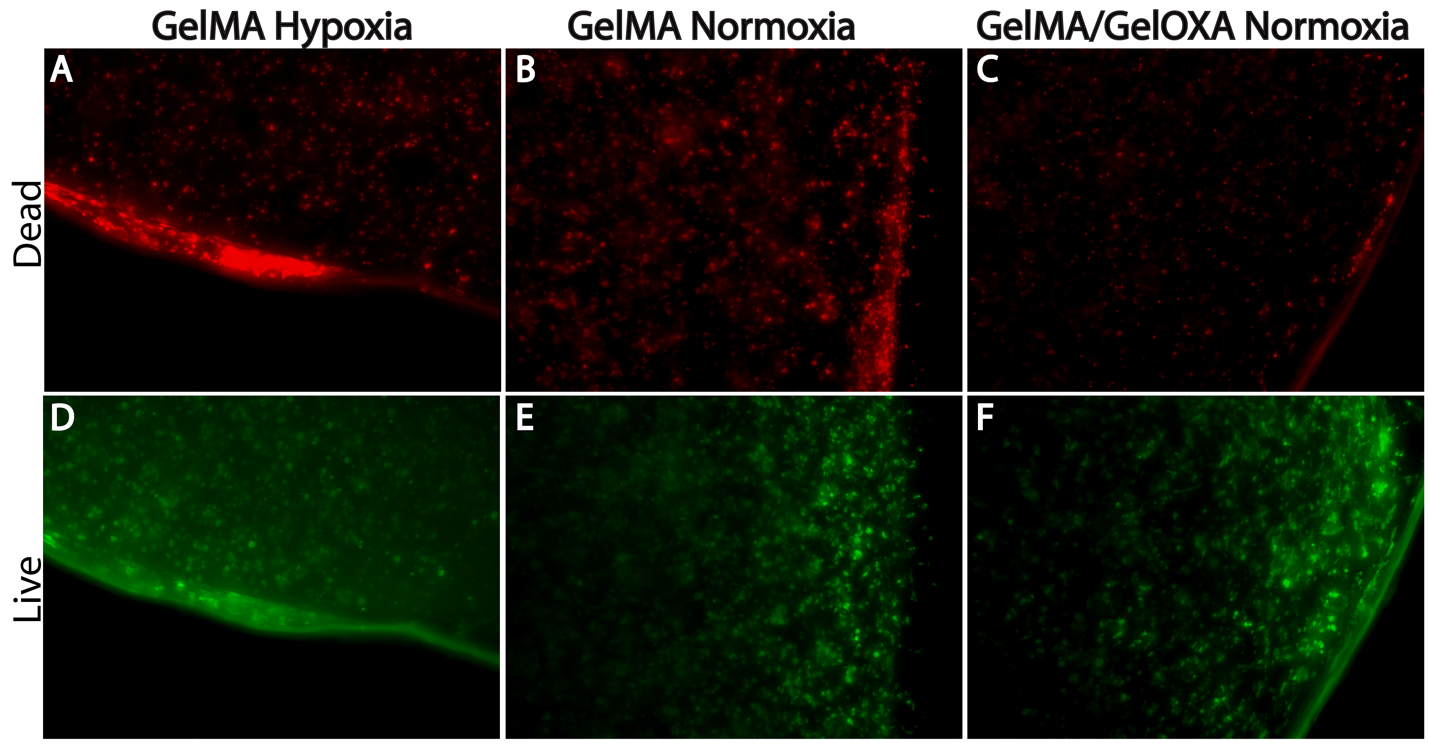

**SI 5:** Live (D-F), and dead (A-C) staining signal reported as single-channel images.

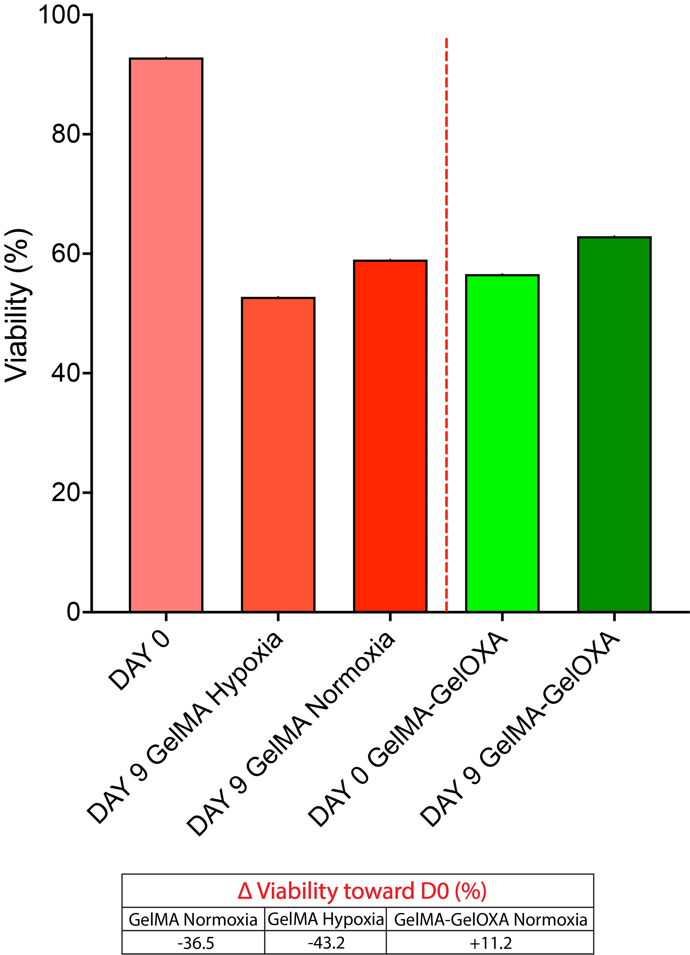

**SI 6:** Quantification of cell viability via live/dead assay performed on cellularized scaffolds.

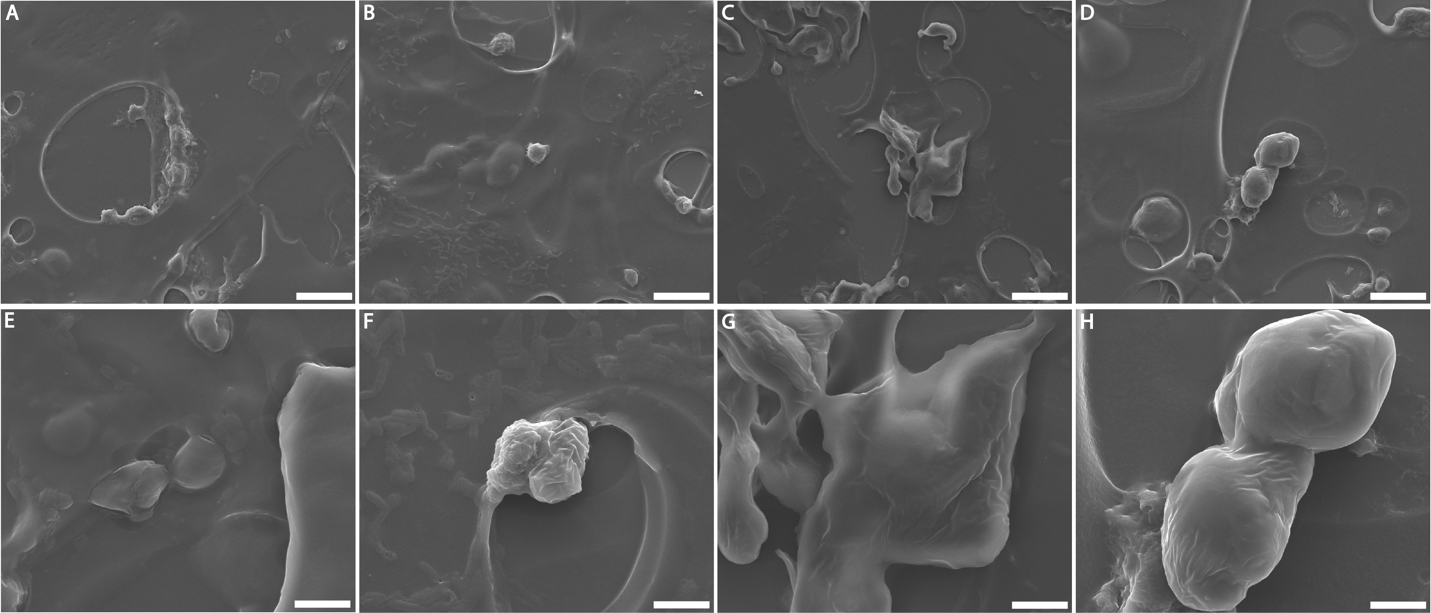

**SI 7:** Images captured by SEM of D0 (A, E), GelMA Hypoxia (B, F), GelMA Normoxia (C, G), and GelMA/GelOXA Normoxia (D, H) samples. Scale bar = 20 µm (A-D). Scale bar = 5 µm (E-H).
